# Extracellular matrix stiffness determines the phenotypic behavior of dedifferentiated melanoma cells through a DDR1/2-dependent YAP mechanotransduction pathway

**DOI:** 10.1101/2024.08.26.609700

**Authors:** Margaux Lecacheur, Ilona Berestjuk, Alexandrine Carminati, Océane Bouvet, Serena Diazzi, Pierric Biber, Christopher Rovera, Marie Irondelle, Frédéric Larbret, Virginie Prod’homme, Christophe A. Girard, Marcel Deckert, Sophie Tartare-Deckert

## Abstract

Extracellular matrix (ECM) stiffening, resulting from increased collagen deposition and cross-linking, is a key biophysical factor of the tumor microenvironment. Cutaneous melanoma is a deadly metastatic cancer. Its aggressiveness stems from high intratumoral heterogeneity, resulting from the plasticity of melanoma cells, which transit from a melanocytic state to dedifferentiated therapy-resistant and invasive phenotypes, characterized by mesenchymal and/or neural crest stem cell-like features. Phenotypic plasticity is regulated by stroma-derived soluble factors, but the functional impact of ECM stiffening on melanoma cell phenotypes remains ill defined. Here, we found that melanoma cell subpopulations display difference in mechanical responsiveness. Compared to melanocytic cells, mesenchymal dedifferentiated cells showed increased proliferation, migration and resistance to MAP kinase-targeted therapy when seeded on stiff collagen. By contrast, a soft ECM impaired their proliferation and migration and sensitized them to targeted therapy. In addition, extracellular mechanical signals are required to sustain melanoma cell identity and dedifferentiation features. Further analyses indicated that the mechanosensitivity nature of dedifferentiated cells relies on the expression and activation of collagen receptors DDR1 and DDR2 that control actomyosin cytoskeleton reorganization and YAP mechanotransduction pathway. Inhibiting both DDR in dedifferentiated melanoma cells abrogated their mechano-induced behavior and drug-resistant phenotype, while forcing their expression in melanocytic cells induced mechanical responsiveness and a less differentiated phenotype. Our results reveal that phenotypic reprogramming endows dedifferentiated melanoma cells with increased sensitivity and addiction to ECM stiffness. We propose that mechano-addiction mediated by DDR collagen receptors may represent a novel vulnerability for aggressive dedifferentiated cancer cells that can be exploited for therapeutic benefits.

## Introduction

Extracellular matrix (ECM) stiffness-induced signaling pathways play a crucial role in cell behavior and tissue homeostasis [1]. The ECM, a complex and dynamic network of proteins, provides structural support to surrounding cells and regulates cellular functions [2]. When the ECM becomes stiffer, cells sense and respond to these mechanical cues through ECM receptors and signaling pathways that influence gene expression [3]. These pathways include integrin-mediated focal adhesion kinase (FAK) activation, Rho/ROCK-actomyosin signaling, and the Hippo-YAP/TAZ pathway [4, 5]. While the role of β1-integrin in mechanotransduction has been well described, the contribution of other ECM receptors such as the discoidin domain tyrosine kinase receptors DDR1 and DDR2 is less well appreciated. Activation of mechanosensitive pathways modulates cell growth, survival, migration and stem cell fate, ultimately influencing tissue development and disease progression [1]. In the context of cancer, altered ECM deposition and increased collagen cross-linking leading to tumor stiffening is known to foster the growth, progression and recurrence of many solid tumors [1].

Melanoma is the leading cause of skin cancer-related death due to its high degree of intratumoral heterogeneity and tumor cell plasticity leading to therapy failure and metastatic propensity. Most mutations found in cutaneous melanoma affect MAP kinase signaling pathway key components such as *BRAF* (40-50%), *NRAS* (20-30%), and *NF1* (10-15%) and lead to its constitutive activation [6]. BRAF-mutated metastatic melanomas are treated with targeted therapy combining BRAF and MEK inhibitors (BRAFi/MEKi) or immune checkpoint inhibitors. BRAFi/MEKi combotherapy is used for patients who require urgent treatment as it provides faster disease control, while immunotherapies are reserved for slow progressing metastatic melanoma as they provide more durable response [7]. While both treatments have improved the clinical outcomes of metastatic melanoma, not all patients respond to these regimens, and those who initially respond eventually relapse due to the development of resistance.

Melanoma is notorious for its intratumor heterogeneity, resulting from the ability of melanoma cells to dynamically oscillate between distinct transcriptional states or phenotypes. This phenotypic plasticity enables them to survive and rapidly adapt to microenvironmental cues and stressful conditions. The most identified melanoma cell states are the melanocytic phenotype and the mesenchymal-like dedifferentiated lineage phenotype [8]. Melanocytic cells are proliferative and drug sensitive. They are differentiated and express high levels of melanocyte lineage specific microphtalmia-associated transcription factor (MITF) and low levels of the receptor tyrosine kinase AXL. In contrast, dedifferentiated cells are poorly proliferative, invasive and drug resistant. They show low MITF expression and high expression of mesenchymal markers including AXL or ZEB1 [9, 10]. Cooperation between these cell states has been shown to promote melanoma progression and metastasis through heterotypic cluster formation [11–13]. Transition from a melanocytic to a dedifferentiated phenotype occurs through multiple intermediate cell states including a neural crest stem-cell like (NCSC) state, which contributes to drug tolerance and tumor relapse [14].

Melanoma cell phenotype switching can be triggered by different cues coming from its microenvironment, including soluble factors (i.e. TGFβ, INFγ), nutrient deprivation, hypoxia or oncogenic BRAF-targeting therapy [15–20]. Besides phenotypic switching, targeted therapy promotes stromal and cancer cell-derived ECM deposition and remodeling, leading to tumor stiffening and drug relapse [21–25]. Notably, physical and structural cues from the ECM confer drug-protective action to melanoma cells against anti-BRAF/MEK therapies [22, 26, 27]. Collagen remodeling is also observed in aging skin and promotes melanoma metastasis [28]. In epithelial cancers, increased ECM stiffness promotes metastasis through the induction of epithelial to mesenchymal transition (EMT), a cell transdifferentiation process resembling melanoma mesenchymal-like transition [29, 30]. More recently, collagen abundance has been shown to promote melanoma cell differentiation, suggesting that ECM rigidity may control phenotypic plasticity [31]. However, it remains to be determined whether ECM stiffness differently impacts on melanoma cell states to regulate their malignant features and aggressiveness.

Here, we sought to investigate how mechanical signals from the ECM influence melanoma cell phenotypic behavior and demonstrated that melanoma cell subpopulations display difference in mechanical responsiveness. We selected a panel of melanoma cell lines and patient-derived melanoma short-term cultures presenting either a melanocytic proliferative or dedifferentiated invasive phenotype and exposed them to collagen-coated substrates of increasing stiffness. We present evidence that lineage dedifferentiation increases sensitivity and responses to stiff ECM and identify the collagen receptors DDR1 and DDR2 as key players in mediating mechano-responses in dedifferentiated melanoma cells. Our work reveals that ECM stiffness differentially impacts on melanoma cell phenotypes and highlight a novel dependency of dedifferentiated melanoma cells on mechanical signals that can be used to target this highly aggressive, treatment-refractory cancer cell state.

## Materials and methods

### Cells and reagents

Melanoma cell lines 501Mel (RRID:CVCL_4633) were obtained as described before [32, 33]. 1205Lu cells (RRID:CVCL_5239) were from Rockland (USA). Short-term patient derived cells were provided by Jean-Christophe Marine [34]. Cells lines were cultured in Dulbecco’s modified Eagle Medium (DMEM) plus 7% FBS (Hyclone) and 1% Penicillin/Streptomycin (Gibco). Short term derived cells were cultured in Ham’s F10 nutrient mix (Invitrogen) with 10% FBS (Hyclone), 7% Glutamax (Gibco) and 1% penicillin/streptomycin (Invitrogen). To guarantee cells authenticity, cells were expanded and frozen at low passage after their receipt from original stocks and used for a limited number of passages after thawing. All cell lines were routinely tested for the absence of mycoplasma by PCR. Culture reagents were purchased from Thermo Fisher Scientific. BRAF inhibitor (Dabrafenib), MEK inhibitor (Trametinib), ROCK inhibitor (Y27632), myosin II inhibitor (Blebbistatin) and DDR inhibitor (DDR1in1) were from Selleckchem. DDR inhibitor (Imatinib mesylate) was from Enzo Life Science. Adenoviruses expressing DDR1 and DDR2 were from Applied Biological Materials, and were amplified as described [33].

### Preparation of substrates of different rigidity

Polyacrylamide hydrogels: Hydrogels were generated from a mixture of 40% acrylamide (Fisher Bioreagents) and 2% bisacrylamide (Fisher Bioreagents) to obtain different rigidities. This mixture was cast between two glass slides. After 10min of polymerization at room temperature, the hydrogels were coated for 45min at 37°C with type I collagen and then sterilized under UV for 30min. After 45min equilibration in DMEM medium, cells can be seeded on it.

Collagen gels: Soft collagen gels were made by mixing bovine type I collagen (2.5 mg/ml) in PBS and NaOH (0.1 N), adjusting the pH of the mixture to approximately 7 as indicated by pH paper. The mixture was added to the glass-bottom section of the dish and incubated for 1h or until completion of the gelation at 37°C. Stiff substrates were generated by coating tissue culture dishes with bovine type I collagen at 50µg/ml for 1h at 37°C.

### Cell proliferation

Cells were plated in 48-well dishes at 10^4^ cells in triplicate on soft or stiff substrate and grown for 3 days. Proliferation was followed for 3 days by counting the nuclei of NucLight Red-expressing cells using the IncuCyte ZOOM™ system according to the manufacturer’s instructions (Essen BioScience). Alternatively, cells were fixed with paraformaldehyde and labeled with Ki67 and Hoechst (33342, Invitrogen). Cells were imaged using a fluorescence microscope (Leica DMI6000, 5X magnification).

### Flow cytometry

#### Cell cycle analysis

Cell cycle profiles were determined by flow cytometry analysis of propidium iodide (PI)-stained cells as described before [32]. Melanoma cells plated on soft or stiff substrate and/or treated with the indicated drugs for 72h were detached, washed and fixed in ice-cold ethanol 70% for at least 24h. Cells were then stained for 15min at RT with 40μg/mL PI (Sigma) in FACS Buffer (1X PBS + 0.5% BSA + 1mM EDTA) supplemented with 100μg/mL RNAse A (Sigma). Cell cycle profiles were collected on FACS Canto II cytometer (Becton Dickinson).

#### Apoptosis analysis

Cell death was evaluated by flow cytometry. Cells were plated on soft or stiff substrate and treated with the indicated drugs for 72h. They were then detached, washed in Annexin-V binding buffer (Miltenyi Biotec) and stained with Annexin-V-FITC and PI for 15min at RT in the dark, according to the manufacturer’s instructions (Miltenyi Biotec). Cell death was analyzed on FACS Canto II cytometer (Becton Dickinson). FACS data were analyzed using FlowJo software.

### Time lapse migration assays

Cells were plated in 24-well dishes at 2x10^4^ cells on soft or stiff substrate for 24h. Cells were stained with Hoechst for 5min, then washed and their migration was assessed by time-lapse live imaging for 24h with one image taken every 10min, using the inverted widefield microscope (DMI6000, Leica, 5X) under 37°C and 5% CO2. Cell displacement was analyzed using the semi-automated plugin of Fiji software (ImageJ) and TrackMate [35].

### Invasion assays

Invasion was assessed by Boyden chamber assay using 8µm pore transwell (Corning) coated overnight with 70µl of Matrigel. Cells were seeded for 7 days on soft or stiff substrate and starved for 24h then dissociated and counted. 150 000 cells were loaded in the upper compartment of the transwell and FBS containing-medium was added in the lower compartment of the transwell. After 24h, migrating cells were fixed and nuclei were stained with Hoechst. Pictures were taken with the inverted widefield microscope (DMI6000, Leica, 10X). Quantification was done with Fiji software (ImageJ).

### RNAi studies

siRNA used in this study are listed in Supplementary Table 1. 50nM siRNA were transfected using Lipofectamine RNAi MAX (Thermo Fisher Scientific), following manufacturer’s instructions.

### Immunoblot analysis and antibodies

Cell lysates were subjected to immunoblot analysis as described before [33] using antibodies listed in Supplementary Table 2.

### Real-time quantitative PCR

Gene expression levels were determined by RT-qPCR as described [32]. Data were presented as relative gene expression according to the ΔΔCt method. Heatmaps depicting fold changes of gene expression were prepared using MeV software. Primer sequences are listed in Supplementary Table 3.

### Traction force microscopy (TFM)

Contractile forces exerted by melanoma cells were assessed by TFM as described [36] using collagen-coated polyacrylamide hydrogels with shear modulus of 0.2kPa or 4kPa containing red fluorescent beads (SoftTrac, Cell Guidance Systems). Cells were plated on fluorescent bead–conjugated gels for 48h. Images were acquired before and after cell removal using a fluorescence microscope (Leica DMI6000, 10X magnification). Tractions exerted by cells were estimated by measuring beads displacement fields, computing corresponding traction fields using Fourier transformation and calculating root-mean-square traction using the particle image velocity (PIV) plugin on ImageJ. The same procedure was performed on a cell-free region to measure baseline noise.

### Immunofluorescence analysis

Cells were grown either on collagen coated coverslips or collagen-coated polyacrylamide/bisacrylamide synthetic hydrogels with defined stiffness for 72h. Following treatment, cells were rinsed, fixed in 4% formaldehyde, and incubated in PBS 0.3% triton 5% Goat Serum in PBS for 1h and then with the dilution of indicated primary antibodies overnight at 4°C. Following incubation with Alexa Fluor-conjugated secondary antibodies (1:200), coverslips and hydrogels were mounted in Prolong antifade (Thermo Fisher Scientific). F-actin was stained with Alexa Fluor594 or Alexa Fluor488 phalloidin (1:200, Thermo Fisher Scientific). Nuclei were stained with Hoechst. Images were captured on a widefield microscope (Leica DM5500B, 40X magnification). Quantifications of cell area, YAP nuclear localization and the number and size of focal adhesions were performed with Image J software.

### Statistical analysis

Unless otherwise stated, all experiments were repeated at least three times and representative data/images are shown. Statistical data analysis was performed using GraphPad Prism software. Unpaired two-tailed Mann-Whitney test was used for statistical comparisons between two groups and Kruskal-Wallis test with Dunn posttests or two-way analysis of variance test with Bonferroni post-tests to compare three or more groups. Histogram plots and curves represent mean ± SD and Violin plots represent median +/- quartiles. P values ≤ 0.05 were considered statistically significant.

## Results

### Dedifferentiated melanoma cells are more sensitive to ECM stiffening than melanocytic cells

Melanoma cells are sensitive to ECM stiffening [31 Girard, 2020 #2226], yet whether their mutation status or transcriptional state influences responsiveness to substrate rigidity remains poorly defined. To test this, we selected melanoma cell lines and shortterm melanoma cultures (designated as MM) harboring either a BRAF or NRAS mutation. qPCR analysis of genes associated with melanoma phenotype switching confirmed the MITF^high^/AXL^low^ melanocytic phenotype of 501Mel, MM057, MM074, MM034, MM001 and MM057 cells associated with the increased expression of MITF, SOX10 or MLANA. In contrast, 1205Lu, MM029, MM099, MM047, and MM165 cells exhibit a MITF^low^/AX^high^ dedifferentiated phenotype associated with the increased expression of AXL, SOX9 and ZEB1 (Table 1, Fig. 1A). To assess their sensitivity to changes in ECM stiffness conditions, cells were plated on collagen-coated polyacrylamide hydrogels of low (<1kPa) versus high (>16kPa) stiffness (Fig. 1B). Culturing dedifferentiated 1205Lu cells for 48h on stiff collagen induced an increase in their spreading and cell area, while it had no significant effect on melanocytic 501Mel cells (Fig. 1C). Similar observations were made with melanocytic versus dedifferentiated short-term MM cells (Fig. 1C and Supplementary Fig. 1A). In line with their increased spreading, dedifferentiated melanoma cells formed more mature focal adhesions than melanocytic cells upon increased matrix rigidity (Fig. 1D and Supplementary Fig. 1B). In addition, they showed activation of YAP-dependent mechanosensing pathway, illustrated by YAP nuclear translocation and increased expression of several of its target genes (AMOTL2, CTGF, CYR61) under high collagen stiffness (Fig. 1E and Supplementary Fig. 1C-E). Interestingly, both BRAF-mutated (1205Lu, MM029, MM099) and NRAS-mutated (MM047, MM165) dedifferentiated melanoma cells were much more sensitive to collagen stiffness than BRAF- or NRAS-mutated melanocytic cells, suggesting that mechano-responsiveness was dictated by tumor cell state rather than mutationnal status. Overall, these data indicate that melanoma cells presenting a dedifferentiated phenotype are more sensitive to extracellular mechanical signals than melanocytic melanoma cells, regardless of their mutation status.

**Figure 1:**
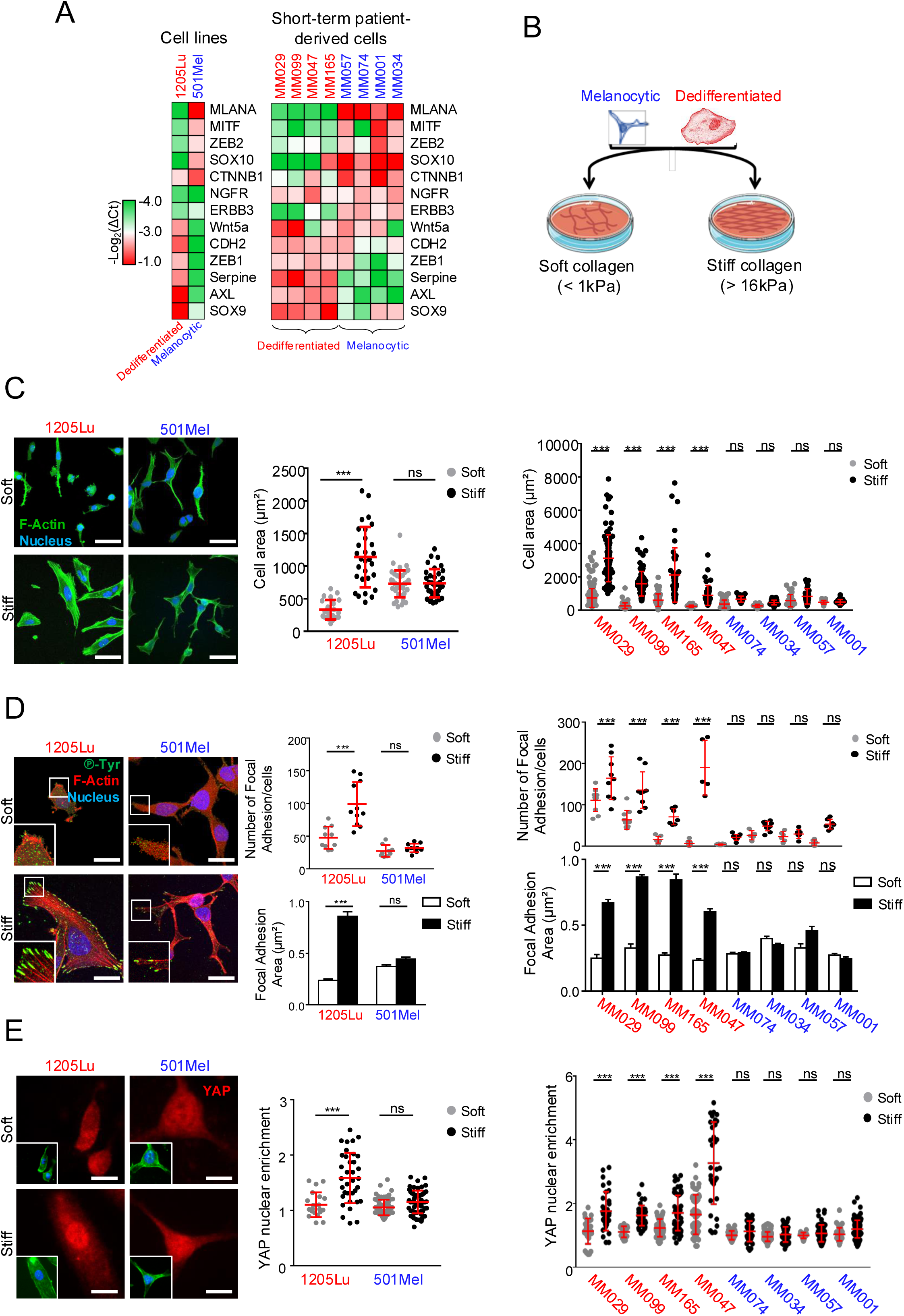
Dedifferentiated melanoma cells display mechanical responsiveness compared to melanocytic cells. (A) Expression heatmap of phenotype switching markers in melanoma cell lines (*left*) and short-term patient derived melanoma cells (*right*) used in the study. Dedifferentiated cells are in red and melanocytic cells in blue. (B) Cartoon illustrating the experimental design used to assess melanoma cell sensitivity to collagen-coated substrates of increasing stiffness. (C) *Left,* Representative images of the morphology of dedifferentiated (1205Lu) or melanocytic (501Mel) melanoma cells cultured for 72h on soft versus stiff collagen-coated substrates. F-Actin was stained with phalloidin (green) and nuclei with Hoechst (blue). Scale bar, 50µm. *Right*, Quantification of morphological changes of melanoma cell lines (*left*) or short-term patient derived melanoma cells (*right*) cultured as above. Data are representative of 3 independent experiments (n > 150 cells). Data are represented as scatter plot with mean ± SD. ***, P<0.001; ns, non-significant, Kruskal-Wallis analysis. (D) *Left,* Immunofluorescence analysis of focal adhesions of 1205Lu or 501Mel cells cultured for 72h on collagen-coated soft or stiff substrate. Focal adhesions were stained with anti-Phospho-Tyrosine antibody (green), F-Actin with phalloidin (red) and nuclei with Hoechst (blue). Insets show higher magnification views of delineated zones. Scale bar, 25µm. *Right-top*, Number of focal adhesions per cell of melanoma cell lines (*left*) or short-term patient derived melanoma cells (*right*). Data are representative of 3 independent experiments. Each dot represents a field with 3-10 cells. (n = 10 fields). Data are represented as scatter plot with mean ± SD. ***, P<0.001; ns, non-significant, Kruskal-Wallis analysis. *Right-bottom*, Bar graphs show the focal adhesion area in cell lines (*left*) or short-term patient derived cells (*right*). Data represent the mean ± SD (n > 150 cells) of triplicate experiments. ***, P<0.001; ns, non-significant, Kruskal-Wallis analysis. (E) *Left*, Effect of collagen stiffening on YAP nuclear translocation assessed by immunofluorescence in cells cultured for 72h on soft or stiff collagen substrates. Insets show F-Actin in green and nuclei in blue. Scale bar, 15µm. *Right*, YAP nuclear enrichment is represented as scatter plot with mean ± SD (n ≥ 30 cells per condition). Data are representative of 3 independent experiments. ***, P<0.001; ns, non-significant, Kruskal-Wallis analysis.

**Table 1:** Phenotypic state and mutation status of the different melanoma cell lines and short-term patient derived melanoma cells used in the study.

|  | Mutation status | Phenotypic state |
| --- | --- | --- |
| *Cell line* |  |  |
| 1205Lu | BRAFV600E | Dedifferentiated |
| 501Mel | BRAFV600E | Melanocytic |
| *Short-term patient-derived cell* |  |  |
| MM029 | BRAFV600E | Dedifferentiated |
| MM099 | BRAFV600E | Dedifferentiated |
| MM074 | BRAFV600E | Melanocytic |
| MM034 | BRAFV600E | Melanocytic |
| MM165 | NRASQ61L | Dedifferentiated |
| MM047 | NRASQ61L | Dedifferentiated |
| MM057 | NRASQ61L | Melanocytic |
| MM001 | NRASQ61L | Melanocytic |

### ECM stiffening increases the proliferation of dedifferentiated melanoma cells and their resistance to BRAF mutant-targeting therapy

ECM-derived mechanical signals have been shown to increase the growth of various solid cancers and to decrease their response to anti-cancer treatments [1, 22, 23, 27]. We therefore examined the functional consequences of collagen stiffening on melanoma cell subpopulations. First, we compared the proliferation of melanocytic and dedifferentiated melanoma cells grown on collagen-coated hydrogels of varying stiffness using IncuCyte live cell imaging. As predicted, melanocytic 501Mel cells were more proliferative that dedifferentiated invasive 1205Lu cells (Fig. 2A). Most melanoma cells studied showed enhanced proliferation in response to increased substrate stiffness in a manner independent of their differentiation status. However, the enhanced proliferation was stronger with dedifferentiated cells when compared to melanocytic cells, with no detectable effect observed for MM057 and MM001 melanocytic cells (Fig. 2A-C). Comparable results were obtained using a Ki67 proliferation marker staining (Supplementary Fig. 2A-B). We also observed that the cell response to stiff collagen was associated with high expression of the differentiation marker SOX10 in 501Mel and MM074 cells, and with high MITF levels in 501Mel cells (Supplementary Fig. 2C). Remarkably, the proliferation of dedifferentiated melanoma cells was blocked on soft substrate (Fig. 2A). Cell cycle analysis further showed that low stiffness induced a G0/G1 cell cycle arrest in dedifferentiated melanoma cells whereas it only had a modest effect in differentiated ones (Fig. 2D and Supplementary Fig. 2D). At the molecular level, this was associated in dedifferentiated cells with a strong reduction in the expression of proliferation markers P-Rb and E2F1 (Fig. 2E) and mesenchymal/resistant markers AXL and ZEB1 (Supplementary Fig. 2C). Altogether, these results suggest that the phenotypic pattern and the proliferation of dedifferentiated melanoma cell are dependent on high ECM stiffness.

**Figure 2:**
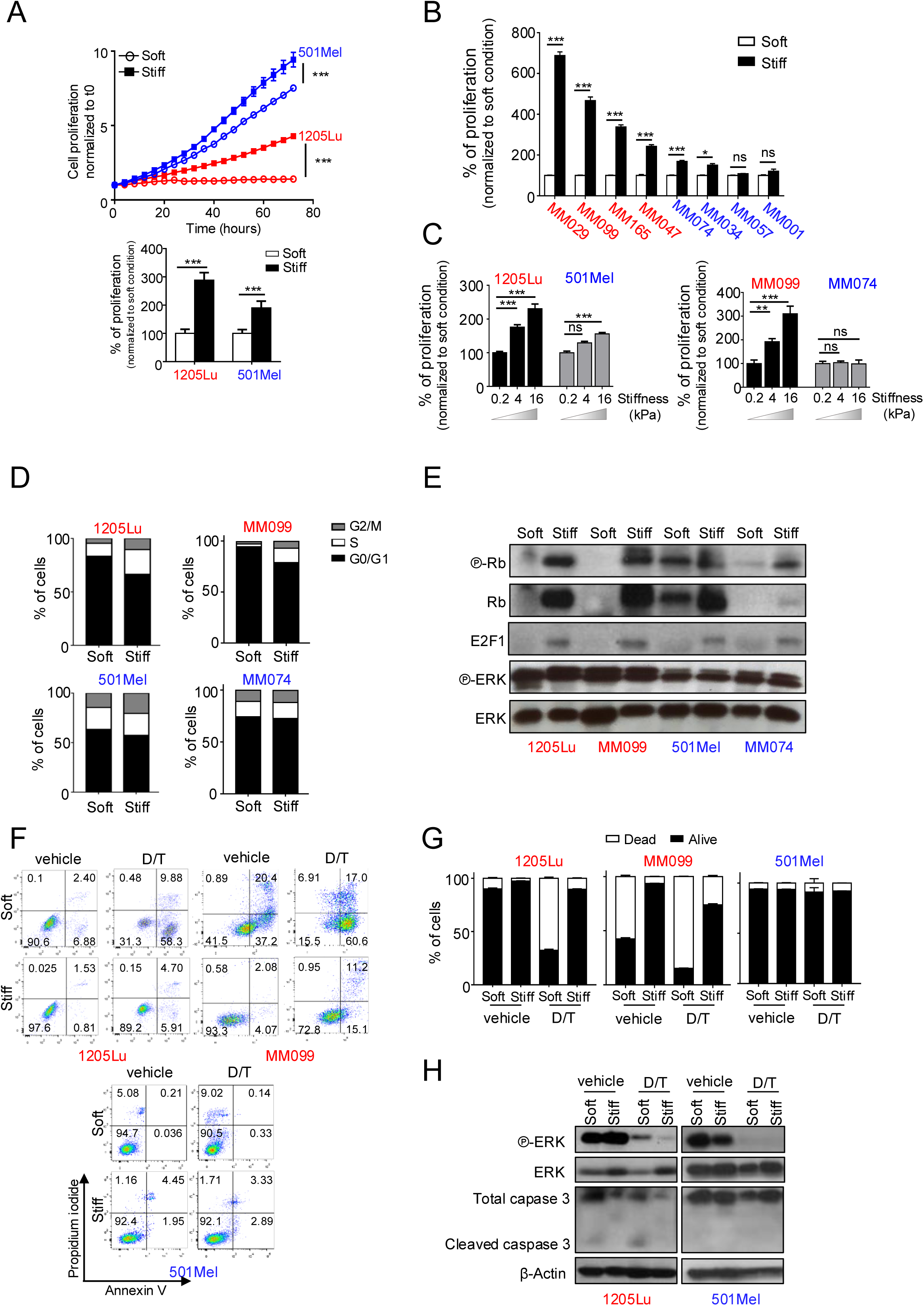
Dedifferentiated melanoma cells display mechano-dependent proliferative and drug resistance properties. (A) *Top,* Time lapse analysis of 501Mel or 1205Lu melanoma cell lines proliferation cultured for 72h on soft or stiff collagen-coated gels, analyzed using the IncuCyte system. Graphs show quantification of cell numbers from NucLight Red nuclear object counting. Data are the mean ± SD, from a minimum of 10 fields per conditions. ***, P<0.001, two-way ANOVA analysis. *Bottom*, Bar graph of cell proliferation after 72h culture on soft or stiff collagen-coated substrate. Data are normalized to proliferation on soft condition ***, P<0.001, Kruskal-Wallis analysis (n = 3). (B) Bar graph of shortterm patient-derived melanoma cell proliferation measured at 72h post seeding on soft or stiff collagen-coated substrate. Data are normalized to proliferation on soft condition ***, P<0.001, Kruskal-Wallis analysis; ns, non-significant, Kruskal-Wallis analysis. Data are representative of 3 independent experiments. (C) Bar graph of proliferation of 1205Lu versus 501Mel cell lines (*left*) or MM099 versus MM074 short-term patient derived cells (*right*) seeded on polyacrylamide collagen-coated hydrogels of increasing stiffness. Data are normalized to proliferation on soft condition. Data are representative of 3 independent experiments. ***, P<0.001; ns, non-significant, Kruskal-Wallis analysis. (D) Cell cycle distribution of melanoma cell lines (1205Lu or 501Mel) or short-term patient-derived melanoma cells (MM099 or MM074) plated for 72h on the substrates as in (A). (E) Immunoblot analysis of cell cycle markers in lysates from cells obtained from (D). ERK is used as loading control. (F) Quantification of apoptosis in 1205Lu, MM099 or 501Mel cells seeded on soft or stiff collagen-coated substrate and exposed for 72h to vehicle or 1µM of Dabrafenib and 0.1µM of Trametinib (D/T). The percentage of apoptotic cells was determined by flow cytometry analysis of annexin V-FITC/PI-stained cells. Early and late apoptotic cells are shown in the right lower quadrant and right upper quadrant, respectively. (G) Bar graph showing the percentage of dead (*white*) and live (*black*) cells from (F). Data represent the mean ± SD of triplicate experiments. (H) Immunoblot analysis of ERK1/2 phosphorylation and Caspase 3 cleavage in lysates from 1205Lu or 501Mel melanoma cell lines obtained from (F). β-Actin is used as loading control.

We next wondered whether the increased mechanosensitivity of dedifferentiated melanoma cells influences their response to oncogenic BRAF inhibition. To address this question, we seeded dedifferentiated (1205Lu, MM099) or melanocytic (501Mel, MM074) cells on soft or stiff collagen and exposed them for 72h to the combotherapy targeting BRAF (BRAFi, Dabrafenib) and MEK (MEKi, Trametinib). When seeded on stiff substrate, Dabrafenib/Trametinib (D/T) combination induced a G0/G1 cell cycle arrest in both melanocytic and dedifferentiated cells (Supplementary Fig. 2E). However, on soft substrate D/T increased the number of apoptotic cells in 1205Lu and MM099 dedifferentiated melanoma cells, but not in 501Mel melanocytic cells (Fig. 2F-G). In line with, we observed the cleavage of caspase 3 only when 1205Lu cells were seeded on soft substrate and exposed to D/T combination (Fig. 2H). As control, phosphorylation of ERK/MAP kinase was strongly reduced upon administration of the drugs in soft and stiff conditions (Fig. 2H). These data demonstrate that low collagen stiffness significantly increases sensitivity to BRAFi/MEKi combination and, conversely, that ECM stiffening confers drug resistance to dedifferentiated melanoma cells. Further, they also indicate that the response of melanocytic differentiated cells to targeted therapy is independent of ECM stiffness.

### ECM stiffening promotes the migratory and invasive behavior of dedifferentiated melanoma cells

We next assessed the effect of ECM stiffness on motility and invasiveness of melanoma cells. First, we seeded 501Mel melanocytic or 1205Lu dedifferentiated cells on collagen substrates with increasing stiffness and monitored their individual migration using live cell imaging. Neither 501Mel nor 1205Lu cells migrated when plated on soft collagen substrate. However, plating cells on stiff collagen induced an increase in 1205Lu cell motility while it only slightly increased 501Mel cell motility (Fig. 3A and C). Similar results were obtained in short-term melanocytic and dedifferentiated melanoma cells (Fig. 3B and C). As cell migration depends on the ability of cells to contract their actomyosin cytoskeleton and the surrounding substrate, we compared traction stresses generated by melanoma cell lines or short-term cultures using TFM. We observed that dedifferentiated cells applied stronger forces on collagen-coated stiff matrices than melanocytic cells (Fig. 3D-E). Finally, we questioned whether priming melanoma cells on different rigidity would affect their invasive properties. To do so, we cultured melanoma cells on soft or stiff collagen substrates for a week, dissociated them and assessed their invasion ability using transwell chambers coated with matrigel (Fig. 3F). We found that pre-culturing dedifferentiated 1205Lu cells on stiff substrate increased their invasion ability (Fig. 3G). In contrast, priming melanocytic 501Mel cells on stiff substrate did not affect their invasion ability. Similar observations were obtained with short-term melanoma cells (Fig. 3H). Together these results suggest that a stiff ECM controls the migratory and invasive phenotypes of dedifferentiated melanoma cells.

**Figure 3:**
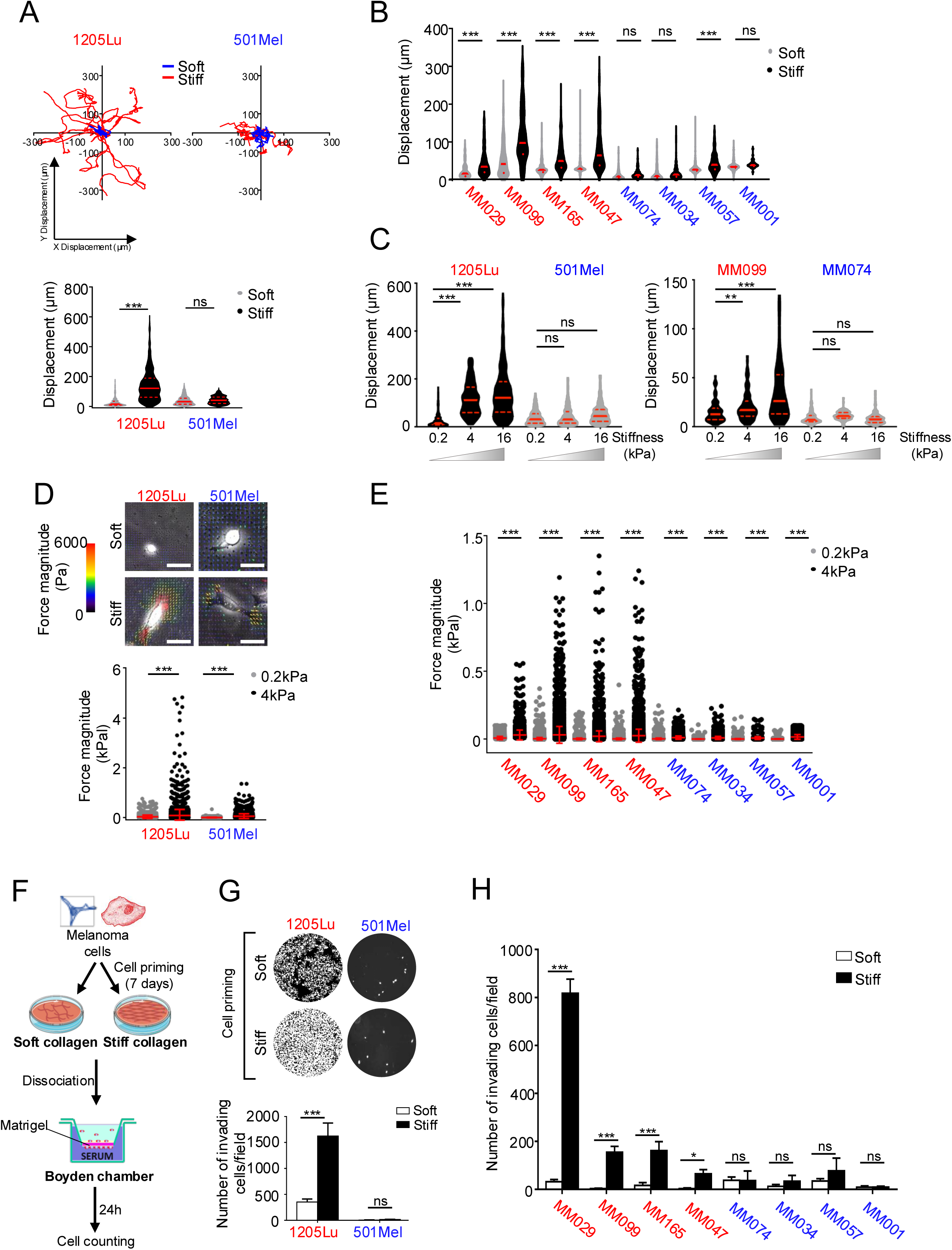
Collagen stiffening increases dedifferentiated melanoma cell motile and invasive behavior. (A) *Top,* Dedifferentiated (1205Lu) or melanocytic (501Mel) melanoma cells were seeded on soft or stiff collagen-coated substrates for 24h, and cell migration was recorded over 24h and automatically tracked using Image J. For each condition, spider plots were created and cell displacement was analyzed using the ImageJ. *Bottom*, Quantification of cell displacement (n = 200 cells). Data represent median ± quartiles. ***, P<0.001; ns, non-significant, Kruskal-Wallis analysis. (B) Violin plot representing cell displacement of short-term patient derived cells seeded as in (A). Data represent median ± quartiles. ***, P<0.001; ns, non-significant, Kruskal-Wallis analysis. (C) Quantification of cell displacement of indicated melanoma cells seeded on collagen-coated polyacrylamide hydrogels of increasing stiffness for 24h (n < 300 cells). Data represent median ± quartiles. Data are representative of 3 independent experiments. ***, P<0.001; ns, non-significant, Kruskal-Wallis analysis. (D) *Top,* Representative images of heat scale plot showing the traction forces applied by 1205Lu or 501Mel cells seeded on 0.2 or 4kPa fluorescent bead-embedded collagen coated hydrogels for 24h. Scale bar, 75µm. *Bottom*, Quantification of contractile forces. Data are the mean ± SD (n = 10 cells/field from 5 random fields). Data are representative of 3 independent experiments. ***, P<0.001; ns, non-significant, Kruskal-Wallis analysis. (E) Scatter plot representing the quantification of contractile forces applied by short-term patient derived melanoma cells on 0.2 or 4kPa fluorescent bead-embedded collagen coated hydrogels for 24h. Data are represented as mean ± SD of 3 independent experiments. ***, P<0.001; ns, non-significant, Kruskal-Wallis analysis. (F) Dedifferentiated or melanocytic melanoma cells were seeded for 7 days on soft or stiff collagen-coated substrates, then detached and counted. Their invasive properties were subsequently analyzed in Boyden chamber for 24h. (G) Representative images (*top*) and quantification (*bottom*) of invading cells in the different conditions. Bar graph represents the mean ± SD of 3 independent experiments. ***, P<0.001; ns, non-significant, Kruskal-Wallis analysis. (H) Quantification of invasive properties of indicated short-term patient derived cells treated as in (F). Data represent the mean ± SD of 3 independent experiments. ***, P<0.001; *, P< 0,05; ns, non-significant, Kruskal-Wallis analysis.

### Collagen receptors DDR1 and DDR2 mediate melanoma cell mechano-responsiveness

Previous studies have shown that members of the integrin and DDR receptors can sense the physical properties of the ECM by interacting with some components of the mechanotransduction pathway [27, 37–39]. Although β1-integrin was overexpressed in dedifferentiated compared to melanocytic melanoma cells, only DDR1 and DDR2 expression and phosphorylation levels appeared modulated by substrate stiffness (Fig. 4A). This led us to investigate the contribution of DDR1 and DDR2 in dedifferentiated melanoma cell mechano-responsiveness. To address this, we either used a siRNA approach to downregulate their expression (Fig. 4B) or a pharmacological approach to inhibit their tyrosine kinase enzymatic activity (Supplementary Fig. 3A). Immunoblot analysis showed that both approaches led to a similar decrease in phosphorylation of myosin light chain 2 (MLC2), the regulatory subunit of myosin II and key regulator of actomyosin contractility. This suggests that DDR mechanical activation controls the actomyosin cytoskeleton in dedifferentiated cells. In addition, the simultaneous knockdown of both DDR or their pharmacological inhibition led to a significant decrease in the number and size of focal adhesions assembled by 1205Lu and MM099 cells plated on stiff substrate (Fig. 4C and Supplementary Fig. 3B). Further, both DDR-targeting approaches led to a significant reduction of YAP nuclear localization (Fig. 4D and Supplementary Fig. 3C) as well as its transcriptional activity (Supplementary Fig. 3D). Finally, exploring publicly available gene expression array from the TCGA, we observed a positive correlation between both DDR1 and DDR2 and YAP expression in human melanoma biopsies (Supplementary Fig. 3E), further emphasizing the close link between YAP and DDR. These data strongly suggest that DDR1 and DDR2 are required to mediate ECM mechanical signals in dedifferentiated melanoma cells.

**Figure 4:**
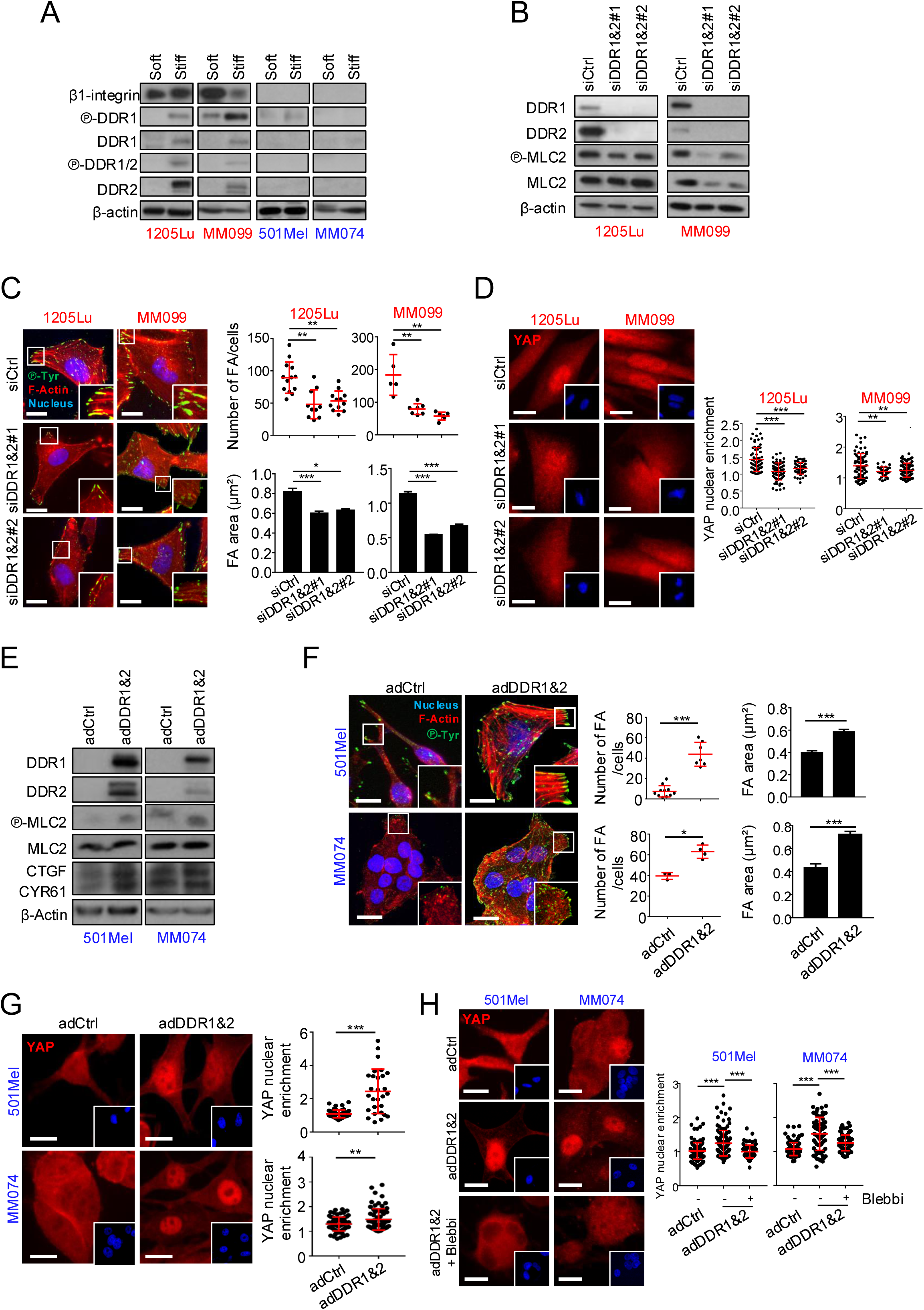
Collagen receptors DDR1 and DDR2 mediate melanoma cell mechano-responsiveness. (A) Immunoblot analysis of β1-integrin, DDR1, DDR2 and MLC2 in lysates from dedifferentiated or melanocytic melanoma cells cultured for 72h on soft or stiff collagen-coated substrate. β-Actin is used as loading control. (B-D) 1205Lu or MM099 dedifferentiated cells on stiff collagen substrate were transfected with the combination of two different sequences of siRNA targeting DDR1 or DDR2 (siDDR1&2#1 and siDDR1&2#2) or control siRNA (siCtrl) (72h, 50nM). (B) Immunoblot analysis of DDR1, DDR2 and MLC2 in lysates from 1205Lu or MM099 cells 72h post transfection with the indicated siRNA. β-Actin is used as loading control. (C) *Left,* Immunofluorescence analysis of focal adhesions (FA) of 1205Lu or MM099 cells described above. FA were stained with anti-Phospho-Tyrosine antibody (green), F-Actin with phalloidin (red) and nuclei with Hoechst (blue). Insets show higher magnification views of zones delineated by a white square. Scale bar, 20µm. *Right*, Quantification of FA per cell (*top*) or FA area (*bottom*). Data are representative of 3 independent experiments. *Top*, Each dot represents a field comprising 3-10 cells. Data are represented as scatter plot with mean ± SD from 10 different fields. ***, P<0.001; ns, non-significant, Kruskal-Wallis analysis. *Bottom*, Bar graph represents the mean ± SD (n > 150 cells) of triplicate experiments. ***, P<0.001; ns, non-significant, Kruskal-Wallis analysis. (D) *Left*, YAP nuclear translocation assessed by immunofluorescence in 1205Lu or MM099 cells described above. Insets show nuclei staining by Hoechst (blue). Scale bar, 20µm. *Right*, Quantification of the nucleocytoplasmic distribution of YAP (n ≥ 30 cells per condition). Data represented mean ± SD of 3 independent experiments. ***, P<0.001; ns, non-significant, Kruskal-Wallis analysis. (E-H) 501Mel or MM074 melanocytic cells seeded on stiff collagen substrate were transduced with the combination of adenoviruses expressing DDR1 or DDR2 (adDDR1&2) or a control adenovirus (adCtrl). (E) Immunoblot analysis of DDR1, DDR2, MLC2 and YAP target genes (CTGF and CYR61). β-Actin is used as loading control. (F) *Left,* Immunofluorescence analysis of focal adhesions (FA) of 501Mel or MM074 cells treated as above. FA were stained with anti-Phospho-Tyrosine antibody (green), F-Actin with phalloidin (red) and nuclei with Hoechst (blue). Insets show higher magnification views of zones delineated by a white square. Scale bar, 20µm. *Right*, Quantification of FA per cell (*top*) or FA area (*bottom*). Data are representative of 3 independent experiments. *Top*, Data are represented as scatter plot with mean ± SD from 10 different fields. Each dot represents a field comprising 3-10 cells. ***, P<0.001; ns, non-significant, Kruskal-Wallis analysis. *Bottom*, Bar graph represents the mean ± SD (n > 150 cells) of triplicate experiments. ***, P<0.001; ns, non-significant, Kruskal-Wallis analysis. (G) *Left*, Immunofluorescence analysis of YAP (red) in 501Mel and MM074 cells. Insets show nuclei staining by Hoechst. Scale bar, 20µm. *Right*, Quantification of the nucleocytoplasmic distribution of YAP (n ≥ 30 cells per condition). Data represented mean ± SD of 3 independent experiments. ***, P<0.001; ns, non-significant, Kruskal-Wallis analysis. (H) *Left*, Representative Images of YAP staining in 501Mel and MM074 cells seeded on stiff substrate and overexpressing DDR1 and DDR2. Cells were treated or not with the myosin II inhibitor Blebbistatin (Blebbi, 40µM) for 72h. Insets represent nuclei stained with Hoechst. Scale bar, 20µm. *Right*, Quantification of the nucleocytoplasmic distribution of YAP (n ≥ 30 cells per condition). Data represent mean ± SD. Data are representative of 3 independent experiments. ***, P<0.001; ns, non-significant, Kruskal-Wallis analysis.

We next asked whether DDR would be sufficient to activate mechanosensitive cell responses. To address this issue, we overexpressed both receptors in melanocytic melanoma cells that exhibit poor mechanoresponsiveness using an adenoviral approach. Interestingly, forcing DDR1 and DDR2 expression in 501Mel or MM074 cells led to an increased phosphorylation of MLC2 (Fig. 4E). DDR-overexpressing melanocytic cells appeared more spread than control cells when exposed to stiff collagen substrate (Supplementary Fig. 3F). This was accompanied by the acquisition of a less differentiated phenotype, as shown by the loss of MITF expression and the increase in ZEB1 expression (Supplementary Fig. 3G). In addition, they formed more mature focal adhesions (Fig. 4F) and displayed an increased nuclear localization of YAP associated with an increase in expression of its target genes (Fig. 4G and Supplementary Fig. 3H). Notably, targeting myosin II with Blebbistatin reversed DDR-induced YAP activation (Fig. 4H). Collectively, these experiments demonstrate that DDR1 and DDR2 control mechanosensitive properties of melanoma cells through the actomyosin-dependent regulation of YAP pathway.

### Targeting DDR, actomyosin or YAP impairs stiff collagen-mediated dedifferentiated melanoma cell phenotypic behavior

We next sought to investigate the consequences of modulating DDR-dependent mechanosignaling on proliferation and migration of melanoma cells. First, we used siRNA to knockdown DDR1 and DDR2 or YAP in 1205Lu or MM099 cells. Supplementary Fig. 4A shows the efficacy of YAP depletion by two different sequences of siRNA in both cell lines. Depleting DDR or YAP reduced cell proliferation (Fig. 5A) and migration (Fig. 5B and C) induced by stiff ECM in dedifferentiated cell states. Consistent with the decreased in MLC2 phosphorylation observed upon DDR knockdown (Fig. 4B), DDR or YAP-silenced dedifferentiated cells generated weaker traction forces on their substrate (Fig. 5D). These results were confirmed by inhibiting DDR tyrosine kinase activities with Imatinib or DDR1-in1 or impairing actomyosin cytoskeleton remodeling with the ROCK inhibitor Y27632 or the myosin II inhibitor Blebbistatin in 1205Lu and MM099 cells (Supplementary Fig. 4B-E). Conversely, ectopic expression of DDR1 and DDR2 was able to trigger high stiffness-dependent migration in differentiated melanocytic cells (Fig. 5E). In addition, 501Mel and MM074 cells overexpressing DDR applied stronger traction forces than control cells when exposed to collagen coated stiff substrate (Fig. 5F). Notably, depleting YAP in those cells prevented mechano-mediated migration (Fig. 5G). These data indicate that loss of rigidity sensing through DDR/actomyosin/YAP targeting suppresses mechanosensitive proliferation and migration in dedifferentiated melanoma cells.

**Figure 5:**
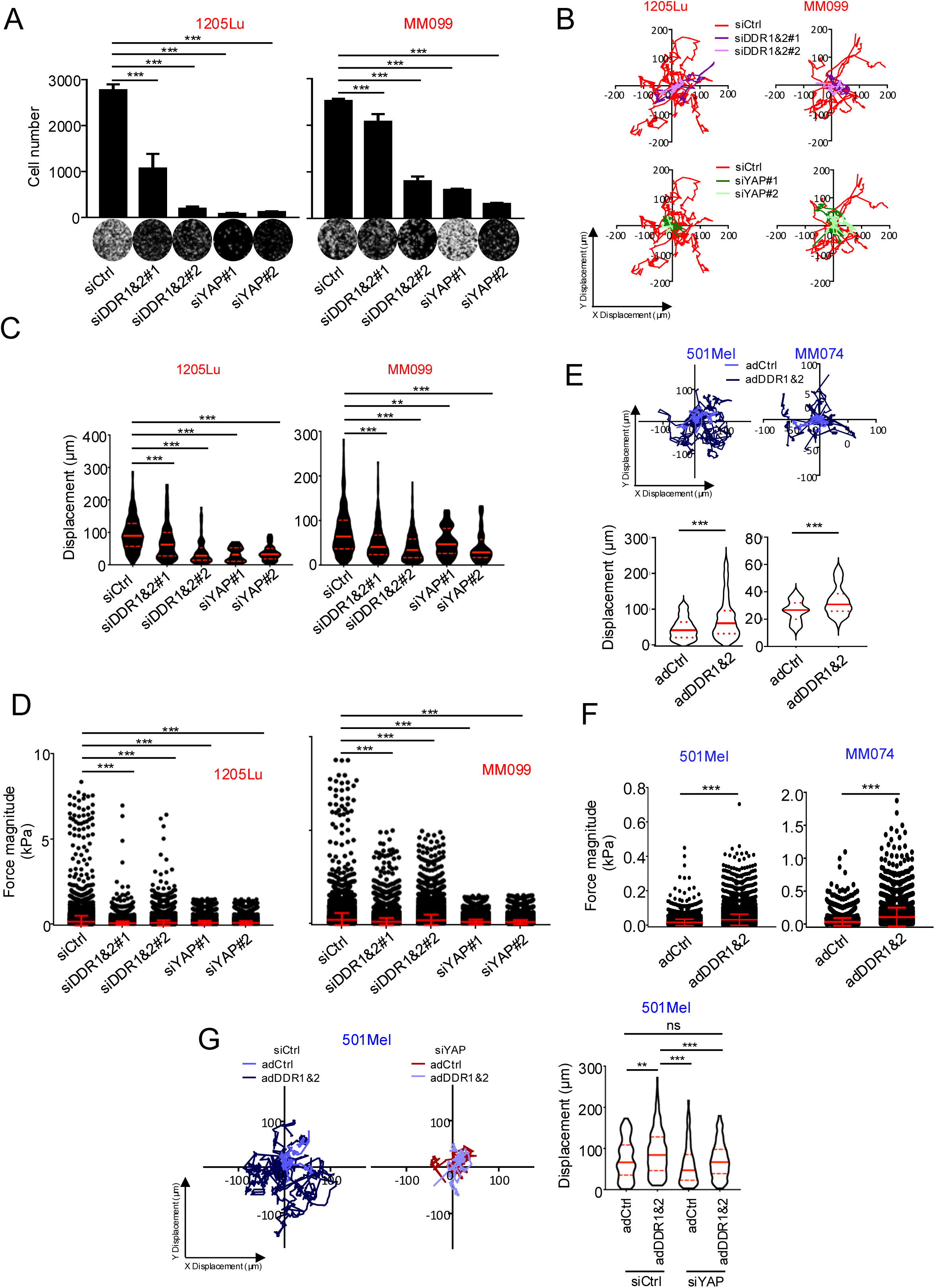
Targeting DDR or YAP impairs ECM stiffness-mediated proliferation and migration in dedifferentiated melanoma cells. (A) Proliferation analysis of 1205Lu and MM099 melanoma cell cultured on collagen-coated stiff substrate, transfected with two different combination of siRNA against DDR1 or DDR2 (siDDR1&2#1 and #2) or against YAP (siYAP#1 and #2). Cell nuclei were stained with Hoechst and counted using ImageJ software. Data are representative of 3 independent experiments. ***, P<0.001; ns, non-significant, Kruskal-Wallis analysis. (B) 1205Lu or MM099 cells treated as in (A) were seeded on stiff collagen for 24h, and cell migration was recorded over 24h and automatically tracked using ImageJ. For each condition, spider plots were created and cell displacement was analyzed using the ImageJ. (C) Quantification of cell displacement (n = 200 cells) from (B). Data represent median ± quartiles. Data are representative of 3 independent experiments. ***, P<0.001; ns, non-significant, Kruskal-Wallis analysis. (D) Quantification of contractile forces applied by 1205Lu or MM099 cells transfected with two combinations of siRNA against DDR1 and DDR2 or YAP for 72h and seeded on 4kPa fluorescent bead-embedded collagen coated hydrogels for 24h. Data are the mean ± SD (n = 10 cells/field from 5 random fields). Data are representative of 3 independent experiments. ***, P<0.001; ns, non-significant, Kruskal-Wallis analysis. (E) *Top,* Cell displacement analysis of 501Mel and MM074 cells transduced with a control (adCtrl) or a combination of DDR1 and DDR2 adenoviruses (adDDR1&2) and seeded on stiff substrate for 24h. *Bottom*, Quantification of cell displacement (n < 300 cells). Data represent median ± quartiles. Data are representative of 3 independent experiments. ***, P<0.001; ns, non-significant, Kruskal-Wallis analysis. (F) 501Mel and MM074 cells transiently overexpressing DDR1 and DDR2 for 72h were seeded on 4kPa fluorescent bead-embedded collagen coated hydrogels for 24h. Data are the mean ± SD (n = 10 cells/field from 5 random fields). Data are representative of 3 independent experiments. ***, P<0.001; ns, non-significant, Kruskal-Wallis analysis. (G) 501Mel cells were transduced with a control (adCtrl) or a combination of DDR1 and DDR2 adenoviruses (adDDR1&2) prior transfection with a control (siCtrl) or a YAP targeting siRNA (siYAP) *Left,* Displacement analysis of 501Mel cells seeded on stiff substrate for 24h. *Right*, Quantification of cell displacement (n < 300 cells). Data represent median ± quartiles. Data are representative of 3 independent experiments. ***, P<0.001; ns, non-significant, Kruskal-Wallis analysis.

### Targeting DDR or YAP sensitizes mechano-addicted melanoma cells to oncogenic BRAF pathway inhibition

Given our previous findings that DDR contribute to ECM-mediated protection against BRAF-targeting therapy in BRAF-mutated melanoma cells [27], we investigated whether DDR are involved in ECM stiffness-induced resistance to the targeted therapy described above. We silenced DDR1 and DDR2 or YAP by siRNA in dedifferentiated cells and evaluated their responses to D/T combination treatment on stiff collagen substrate. Apoptotic cell death was monitored by flow cytometry using annexin V-FITC/PI staining assay and the analysis of the subG1 population. Knockdown of DDR or YAP in 1205Lu or MM099 cells sensitized them to targeted therapy, as illustrated by the increased proportion of cells entering apoptosis (Fig. 6A-B and Supplementary Fig. 5A), and the increased cleavage of caspase 3 observed by immunoblotting (Fig. 6C). Similar observations were obtained after pharmacological inhibition of DDR with Imatinib or DDR1-in1 as well as with the ROCK inhibitor Y27632 or Blebbistatin (Fig. 6D and Supplementary Fig. 5B). Taken together, these results demonstrate that collagen receptors DDR contribute to the drug resistant melanoma cell state observed in response to mechanical signals via the regulation of the actomyosin/YAP pathway.

**Figure 6:**
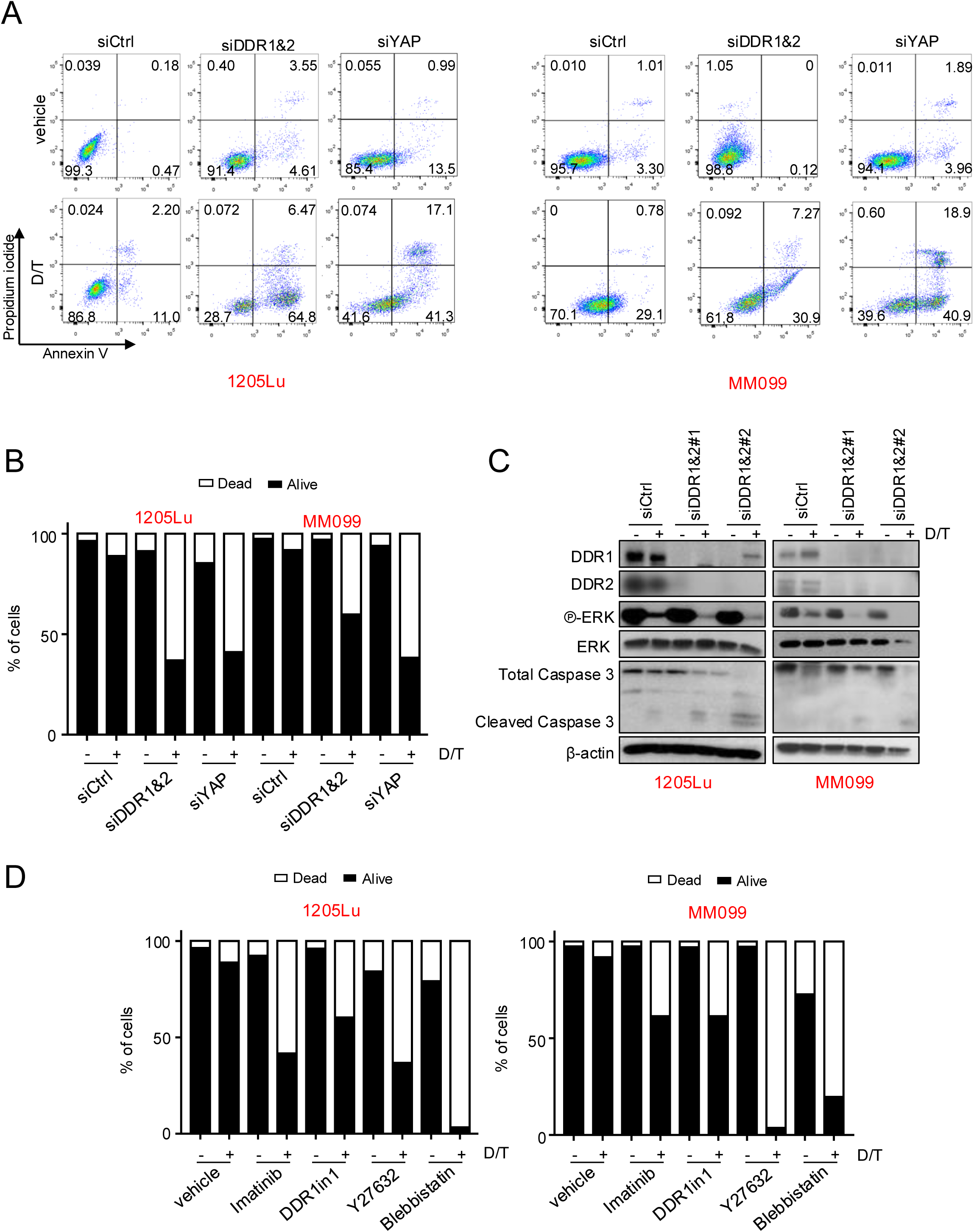
Targeting DDR or YAP sensitizes mechano-addicted melanoma cells to oncogenic BRAF pathway inhibition. (A) Quantification of apoptosis in 1205Lu or MM099 cells seeded on stiff substrate and transfected with siRNA against DDR1 and DDR2 (siDDR1&2) or against YAP (siYAP) prior exposure to vehicle or 1µM of Dabrafenib and 0.1µM of Trametinib combotherapy (D/T) for 72h. The percentage of apoptotic cells was determined by flow cytometry analysis of annexin V-FITC/PI-stained cells. Early and late apoptotic cells are shown in the right lower quadrant and right upper quadrant, respectively. (B) Bar graph showing the percentage of dead (*white*) and live (*black*) cells from (A). Data are representative of 3 independent experiments. Data represent mean ± SD. (C) Immunoblot analysis of ERK1/2 phosphorylation and Caspase 3 cleavage in lysates from 1205Lu and MM099 melanoma cells treated as in (A). β-Actin is used as loading control. (D) Bar graph showing the percentage of dead (*white*) and live (*black*) cells from 1205Lu and MM099 cells seeded on stiff substrate and treated with DDR inhibitors (Imatinib or DDR1in1) or with the ROCK inhibitor Y27632 or the myosin II inhibitor Blebbistatin in combination with 1µM of Dabrafenib and 0.1µM of Trametinib (D/T) for 72h. The percentage of live and dead cells was determined by flow cytometry analysis of annexin V-FITC/PI-stained cells. Data are representative of 3 independent experiments. Data represent mean ± SD.

## Discussion

ECM stiffening is a biophysical hallmark of solid tumors and the result of altered deposition of ECM components and increased collagen cross-linking. It is established that a stiff and fibrotic tumor microenvironment is correlated with increased tumor aggressiveness and poor clinical outcomes in various cancers such as breast and pancreatic carcinomas. In cutaneous melanoma, recent observations have shown that high collagen expression correlates with poorer patient survival [31] and increased fibrous ECM in patients with relapsed melanoma after failure of BRAF inhibitor treatment [23].

Consistently, enhanced fibrillar collagen associated with a pro-fibrotic response and tumor stiffening has been reported during adaptive response and resistance to BRAF-targeted therapy in several pre-clinical mice models [22–25]. Despite the clinical relevance of a mechanical microenvironment in melanoma, little is known about how ECM mechanical signals influence melanoma cell plasticity and phenotypic behavior.

Collagen-coated polyacrylamide hydrogels with variable stiffness is a widely accepted experimental approach for mimicking the matrix environment experienced by tumor cells and studying the functional consequences of changes in ECM stiffness on cellular behavior. Here we report that the transcriptional state of melanoma cells, rather than its mutational status, is the primary determinant of its sensitivity to mechanical cues from the ECM. We show that lineage dedifferentiation towards a mesenchymal-like phenotype determines the sensitivity and responses of melanoma cells to stiff ECM. This data indicates that in contrast to more differentiated melanocytic melanoma cells dedifferentiated melanoma cells empower cellular responses to the mechanical properties of their local microenvironment. This mechano-responsiveness is mediated by the collagen receptors DDR1 and DDR2, which influence phenotypic cell behavior and drug sensitivity. DDR-high dedifferentiated mesenchymal-like melanoma cells showed increased proliferation, migration, and resistance to mutant BRAF-targeted therapy when exposed to stiff collagen. Conversely, melanocytic cells were less responsive to mechanical stiffness, as the result of lower DDR expression [27]. These findings are consistent with data showing that ECM stiffening fosters the aggressiveness of various cancers [29, 37, 40, 41]. In addition to different levels of DDR, the differing responses between melanocytic and dedifferentiated melanoma cells may also be due to variations in TGFβ signaling, which have been reported between these cell types [34]. Miskolczi et al. found that collagen stiffening promotes melanoma cell differentiation and MITF expression in the absence of TGFβ, whereas in the presence of TGFβ, it drives melanoma cell dedifferentiation and upregulates AXL expression [31]. However, our study found that collagen stiffening does not drive the phenotypic switch from a melanocytic to a dedifferentiated state. Instead, it contributes to sustain the existing phenotype by maintaining the expression of differentiation markers like MITF or SOX10 in melanocytic cells, and mesenchymal and drug resistant markers such as AXL, ZEB1 or DDR in dedifferentiated cells. These findings may also explain the coexistence of melanocytic and dedifferentiated melanoma cells in collagen-rich tumor biopsies [31].

We hypothesize that maintaining intratumoral heterogeneity in collagen-dense regions could facilitate cooperation between different melanoma cell subpopulations, a process known to promote invasion and metastasis [11–13]. On the other hand, soft collagen or DDR inhibition blocks melanoma cell proliferation, suggesting that a soft ECM or low DDR signaling may promote the conversion of proliferative cells to slow-cycling or dormant cells. Interestingly, tumor-derived type III collagen has been shown to sustain liver tumor dormancy by blocking DDR1/STAT1-mediated tumor cell proliferation [42]. Whether a similar mechanism operates in melanoma cells within a soft type I collagen environment remains to be determined. An intriguing question is why cells grown on soft collagen are more sensitive to targeted therapies if they are dormant. One possibility is that reducing mitochondrial metabolism upon low DDR mechanical signaling could play a role, supported by the observation that reducing DDR1-dependent adhesion favors the transition from mesenchymal-like melanoma cells with high ATP levels to amoeboid invasive cells with low ATP levels [43].

Mechanistically, DDR1 and DDR2 were identified as key mediators of mechanosensitivity in melanoma cells. Our research, along with that of others, has shown that DDR1 and DDR2 levels increase during melanoma progression and are preferentially enriched in dedifferentiated melanoma cell subpopulations [27]. Targeting DDR1 in human melanoma cell lines decreases their proliferation, invasion, and survival [44, 45], while depleting DDR2 in murine melanoma cell lines reduces their metastatic potential [46]. In line with our findings that both receptors are necessary to mediate fibroblast-derived matrix protection against MAP kinase inhibitors [27], we demonstrated that DDR1 and DDR2 are required to transduce ECM mechanical signals and drug resistance in dedifferentiated cells, while forced expression of DDR in melanocytic cells decreases the expression of the differentiation marker MITF and enhances their response to mechanical cues. Our findings are consistent with studies showing that DDR1 senses matrix stiffness and regulates vascular smooth muscle cells (VSMCs) through RhoA GTPase activation and YAP signaling, creating a feedback loop that further enhances DDR expression [5, 47]. Whether these events involve a direct DDR-myosin II interaction as described for DDR1 during cell migration and collagen mechanical remodeling [39, 48], remains to be determined. However, our data indicate that inhibiting actomyosin remodeling by Blebbistatin impedes DDR-mediated YAP activation of stiff ECM.

Beyond DDR, β1-integrins, another class of collagen receptors play a significant role in mechanosensing [37, 40]. In melanoma, β1-integrins contribute to matrix-mediated resistance to MAP kinase inhibitors [21, 23]. While β1-integrin expression is higher in dedifferentiated cells, it does not change with collagen stiffening as DDR do. However, stiff collagen does increase focal adhesion formation and number, suggesting a role for β1-integrin in supporting the mechanosensitivity of dedifferentiated melanoma cells. The interplay between DDR and β1-integrin may be crucial in conferring these properties, as DDR have been shown to regulate integrin interaction with fibrillar collagen in other cell types [49]. Similarly, DDR2 controls breast tumor stiffness and metastasis by regulating integrin-mediated mechanotransduction in cancer-associated fibroblasts [38].

Our findings provide evidence that stiff collagen enhances drug resistance through the activation of DDR receptors and YAP-dependent mechanotransduction pathways in dedifferentiated melanoma cells. In collagen-rich regions, the increased activity of DDR and YAP in dedifferentiated melanoma cells is likely to exacerbate tumor stroma stiffening, as both DDR and YAP have been implicated in ECM remodeling through cytoskeleton contraction and the production of ECM proteins [39, 50–52]. Notably, our results show that inhibiting DDR or YAP as well as ROCK-Myosin II signaling in dedifferentiated melanoma cells dampens the protective effect conferred by stiff collagen, thereby sensitizing these cells to combined BRAF and MEK inhibition. AXL was shown to be a major player in resistance to targeted therapy in dedifferentiated melanoma cells [9, 53]. Thus, the decrease of AXL on soft ECM is likely to contribute to increased drug sensitivity observed in dedifferentiated cells in this setting. This supports the notion that therapeutic strategies aimed at normalizing tumor-associated fibrosis [24], disrupting melanoma cell-ECM interactions [27], or inhibiting tumor stiffness-induced mechanotransduction pathways [22, 54] could prevent melanoma adaptation to BRAF-targeting therapies. Interestingly, DDR2 inhibition decreases AXL expression and cell proliferation in dedifferentiated BRAFi-resistant melanoma cells, indicating that targeting DDR also counteracts acquired resistance [55].

In addition to resistance to targeted therapies, collagen stiffening, by maintaining the dedifferentiated phenotype of melanoma cells and activating mechanotransduction pathways, may also affect melanoma response to immunotherapies. This idea is supported by evidence linking innate immune checkpoint inhibitor resistance to melanoma subtypes with invasive and dedifferentiated gene expression signatures [56]. Consistently, the up regulation of the EMT transcription factor ZEB1 on stiff collagen may favor melanoma immune evasion and efficacy of immunotherapy [57]. Acquired resistance to anti-MAPK targeted therapies not only promotes an immune-evasive tumor microenvironment but also leads to cross-resistance to immunotherapy in melanoma [58]. This resistance is often associated with highly recurrent non-genomic alterations and shifts in the tumor’s immune landscape, which may contribute to cross-resistance to PD-1/PD-L1 checkpoint inhibitors [16]. ROCK-myosin II signaling further supports the survival of resistant melanoma cells and enhances immunosuppression [54]. Notably, reversing tumor stiffening by inhibiting collagen crosslinking has been shown to improve T cell migration and the efficacy of anti-PD1 treatments [59]. Moreover, DDR2 has shown promise in enhancing tumor responses to anti-PD1 immuno-therapy across several solid cancers, including melanoma [60].

In conclusion, targeting mechanical stimuli from the ECM holds significant potential for enhancing the efficacy of current therapies in melanoma, as well as other collagen-rich and stiff tumors. A deeper understanding of the mechanisms by which ECM stiffness and mechanotransduction contribute to tumor cell behavior and plasticity could pave the way for novel strategies to combat resistant and metastatic cancer cell phenotypes. In this work, we describe differential mechanical responsiveness between different tumor cell phenotypes and demonstrate that mechanical addiction driven by DDR collagen receptors may represent a novel therapeutic vulnerability of the dedifferentiated mesenchymal-like cell state in melanoma.

## Acknowledgments

We thank J.-C. Marine and G.E. Ghanem for providing short-term cultures of melanoma cells. This work was supported by Institut National de la Santé et de la Recherche Médicale (Inserm), Université Côte d’Azur, Ligue Contre le Cancer (Equipe labellisée 2022 to S.T.-D.), Canceropôle Provence Alpes Côte d’Azur (Emergence grant to C.A.G.), Fondation ARC (Projet Fondation ARC to S.T.D.), and Institut National du Cancer (INCA_16697 to S.T.-D.), the French Society of Dermatology (grant to M.D.), ITMO Cancer Aviesan within the framework of the Cancer Plan and the French Government through the ‘’Investments for the Future’’ LABEX SIGNALIFE (ANR-11-LABX-0028-01). We acknowledge the Imaging, Cytometry and Genomics Core Facilities of C3M and the financial contribution of Conseil Départemental 06, Région Sud and Cancéropole Provence Alpes Côte d’Azur. We also thank the “Microscopie Imagerie Côte d’Azur” (MICA) GIS-IBISA platform supported by the GIS IBiSA, Conseil Départemental 06 and Région Sud. M.L., P.B. and O.B are recipients of a doctoral fellowship from Fondation ARC. S.D. is a recipient of a doctoral fellowship from Fondation pour la Recherche Médicale (FRM). A.C. and I.B are recipients of a doctoral fellowship from La Ligue Contre le Cancer.

## Author contributions

C.A.G., M.D. and S.T.-D. designed the study. M.D. and S.T.-D conceptualized and supervised the study. M.L. performed the experiments and analyzed the data with the help of C.A.G., I.B., A.C., S.D., O.B., P.B., C.R., F.L. and V.P. M.I. provided technical advice and expertise in microscopy. C.A.G. and M.L. wrote the original draft. C.A.G., M.L., M.D. and S.T.-D reviewed and edited the final version of the manuscript.

