## Supplemental files for "Extracellular matrix stiffness determines the phenotypic behavior of dedifferentiated melanoma cells through a DDR1/2-dependent YAP mechanotransduction pathway"

#### Supplementary Material

##### Supplementary Figure legends

**Supplementary Figure 1:** (A) Representative images of the morphology of short-term patient derived cells presenting either a melanocytic (in blue) or a dedifferentiated (in red) phenotype cultured for 72h on collagen-coated soft or stiff substrate. F-Actin was stained with phalloidin (green) and nuclei with Hoechst (blue). Scale bar, 50µm. (B) Immunofluorescence analysis of focal adhesions in short-term patient derived cells cultured as in (A). Focal adhesions were stained with anti-Phospho-Tyrosine antibody (green), F-Actin with phalloidin (Red) and nuclei with Hoechst (Blue). Insets show higher magnification views of delineated zones. Scale bar, 20µm. (C) Effect of collagen stiffening on YAP nuclear translocation assessed by immunofluorescence in short-term patient derived cells cultured as in (A). Insets show F-Actin in green and nuclei in blue. Scale bar, 15µm. (D-E) qPCR analysis of YAP target genes expression in melanoma cell lines (D) and short-term patient derived cells (E) cultured as in (A) for 72h. Data are normalized to the expression in cells seeded on soft substrate. Data are represented as mean ± SD from a technical triplicate representative of 3 independent experiments. ***, $P<0.001$; ns, non-significant, two-way ANOVA analysis.

**Supplementary Figure 2:** (A) *Left,* Proliferation of 1205Lu or 501Mel cells cultured for 72h on soft or stiff collagen substrates assessed by Ki67 staining (green). Scale bar, 150µm. *Right*, Bar graph representing the % of Ki67 positive cells per field. 5 fields containing each 20-100 cells were analyzed. Data are representative of 3 independent experiments. Data represent mean ± SD. ***, $P<0.001$; *, $P<0.05$, Kruskal-Wallis analysis. (B) *Top*, Proliferation of indicated short-term patient derived melanoma cells cultured for 72h on soft or stiff collagen-coated substrates, assessed by Ki67 staining (green). Scale bar, 150µm. *Bottom*, Bar graph representing the % of Ki67 positive cells per field. 6-7 fields containing each 20-100 cells were analyzed. Data are representative of 3 independent experiments. Data represent mean ± SD. ***, $P<0.001$; **, $P<0.01$; ns, non-significant, Kruskal-Wallis analysis. (C) Immunoblot analysis of the indicated markers of phenotype switching in lysates from cell lines or short-term patient derived cells cultured for 72h on soft or stiff collagen-coated substrates. HSP90 is used as loading control. (D) Cell cycle distribution of melanoma short-term patient derived cells cultured for 72h on soft or stiff collagen, analyzed by flow cytometry. (E) Cell cycle distribution of 1205Lu, MM099 or 501Mel cells seeded on soft or stiff collagen and treated or not (vehicle) for 72h with 1µM of Dabrafenib and 0.1µM of Trametinib (D/T).

**Supplementary Figure 3:** (A-C) 1205Lu or MM099 dedifferentiated melanoma cells seeded on stiff collagen substrate were treated with vehicle or DDR inhibitors Imatinib (7µM) or DDR1in1 (1µM) for 72h. (A) Immunoblot analysis of activated (phosphorylated) and total forms of DDR1, DDR2 and MLC2 in 1205Lu and MM099 cell lysates. β-Actin is used as loading control. (B) *Left,* Immunofluorescence analysis of focal adhesions (FA) in 1205Lu or MM099 cells. FA were stained with anti-Phospho-Tyrosine antibody (green), F-Actin with phalloidin (red) and nuclei with Hoechst (blue). Insets show higher magnification views of zones delineated by a white square. Scale bar, 20µm. *Right*, Quantification of FA per cell (*top*) or FA area (*bottom*). Data are representative of 3 independent experiments. *Top*, Each dot represents a field comprising 3-10 cells. Data are represented as scatter plot with mean ± SD from 10 different fields. ***, P<0.001; ns, non-significant, Kruskal-Wallis analysis. *Bottom*, Bar graph represents the mean ± SD (n > 150 cells) of triplicate experiments. ***, P<0.001; ns, non-significant, Kruskal-Wallis analysis. (C) *Left*, Immunofluorescence analysis of YAP (red) in 1205Lu and MM099 cells. Insets show nuclei staining by Hoechst. Scale bar, 15µm. *Right*, Quantification of the nucleocytoplasmic distribution of YAP (n ≥ 30 cells per condition). Data represented mean ± SD of 3 independent experiments. ***, P<0.001; ns, non-significant, Kruskal-Wallis analysis. (D) qPCR analysis of YAP target genes expression in 1205Lu or MM099 dedifferentiated melanoma cells seeded on stiff collagen and either transfected with two combinations of siRNA against DDR1 and DDR2 (*top*) or treated with DDR inhibitors (*bottom*) for 72h. Data are normalized to the expression in cells transfected with siCtrl or treated with vehicle, respectively. Data are represented as mean ± SD from a technical triplicate representative of 3 independent experiments. ***, P<0.001; ns, non-significant, two-way ANOVA analysis. (E) Spearman's correlation analysis of DDR1 and YAP (*left*) or DDR2 and YAP (*right*) mRNA expression from data of skin melanoma extracted from the TCGA. (F) *Left,*

Representative Images of 501Mel and MM074 cells cultured on stiff collagen and transduced for 72h with adenoviruses expressing DDR1 and DDR2. F-Actin was stained with phalloidin (green) and nuclei with Hoechst (blue). Scale bar, 50µm. *Right*, Quantification of cell area represented as scatter plot with mean ± SD (n > 150 cells). Data are representative of 3 independent experiments. ***, P<0.001, Kruskal-Wallis analysis. (G) Immunoblot analysis of DDR1, DDR2 MITF and ZEB1 in cell lysates from 501Mel and MM074 cells cultured on stiff substrate and transduced for 72h with adenoviruses expressing DDR1 and DDR2. β-Actin is used as loading control. (H) qPCR analysis of YAP target genes expression in 501Mel and MM074 melanocytic cells transiently overexpressing DDR1 and DDR2 and treated with Blebbistatin (40µM) for 72h on stiff substrate. Data are normalized to the expression in cells infected with control adenovirus (adCtrl) and treated with vehicle. Data are represented as mean ± SD from a technical triplicate representative of 3 independent experiments. ***, P<0.001; ns, non-significant, two-way ANOVA analysis.

**Supplementary Figure 4:** (A) Immunoblot analysis of YAP in cell lysates from 1205Lu and MM099 melanoma cells 72h post-transfection with two different sequences of siRNA against YAP (siYAP#1 and #2). β-Actin is used as loading control. (B) Proliferation analysis of 1205Lu and MM099 melanoma cell cultured on collagen-coated stiff substrate and treated or not (vehicle) for 72h with DDR inhibitors Imatinib (7µM) or DDR1in1 (1µM), ROCK inhibitor Y27632 (40µM) or myosin II inhibitor Blebbistatin (40µM). Cell nuclei were stained with Hoechst and counted using ImageJ software. Data are representative of 3 independent experiments. ***, P<0.001; ns, non-significant, Kruskal-Wallis analysis. (C) *Top*, Cell displacement analysis of 1205Lu and MM099 cells treated with as in (B) and seeded on stiff substrate for 24h. *Bottom*, Quantification of cell displacement (n = 200 cells). Data represent median ± quartiles. Data are representative of 3 independent experiments. ***, P<0.001; ns, non-significant, Kruskal-Wallis analysis. (D) Quantification of contractile forces applied by 1205Lu or MM099 cells treated as in (B) and seeded on 4kPa fluorescent beads-embedded collagen coated hydrogels for 24h. Data are the mean ± SD (n = 10 cells/field from 5 random fields). Data are representative of 3 independent experiments. ***, P<0.001; ns, non-significant, Kruskal-Wallis analysis. (E) Immunoblot analysis of total and phosphorylated forms of MLC2 in cell lysates from 1205Lu and MM099 melanoma cells treated for 72h with the ROCK inhibitor Y27632 (40µM) or myosin II inhibitor Blebbistatin (40µM). β-Actin is used as loading control.

**Supplementary Figure 5:** (A) Cell cycle distribution including sub-G1 phase of 1205Lu and MM099 seeded on stiff collagen and transfected with two combinations of siRNA against DDR1 and DDR2 (siDDR1&2#1 and #2) or siCtrl. (B) Cell cycle distribution including sub-G1 phase of 1205Lu and MM099 seeded on stiff collagen and treated or not (vehicle) with DDR inhibitors (Imatinib or DDR1in1) for 72h.

**Supplementary Table 1**: List of siRNAs

| Name | Target RNA | Reference | Manufacturer |
|---|---|---|---|
| siCtrl | Luciferase | 12935146 | Thermo Fisher Scientific |
| siYAP#1 | YAP1 | #HSS115942 | Thermo Fisher Scientific |
| siYAP#2 | YAP1 | #HSS115944 | Thermo Fisher Scientific |
| siDDR1&2#1 | DDR1 & DDR2 | #HSS187878 & #HSS107350 | Thermo Fisher Scientific |
| siDDR1&2#2 | DDR1 & DDR2 | #HSS187879 & #HSS107352 | Thermo Fisher Scientific |

**Supplementary Table 2:** List of antibodies

| Name | Manufacturer, Ref, RRID | Dilution | Application |
|---|---|---|---|
| AXL | Cell Signaling Technology<br>Cat# 4566,<br>RRID: AB_2062563 | 1:1000 | WB |
| β-Actin | Cell Signaling Technology<br>Cat# 3700,<br>RRID: AB_2242334 | 1:5000 | WB |
| β1-Integrin | Sigma Cat# MAB1965,<br>RRID: AB_2128061 | 1:1000 | WB |
| Caspase 3 (Total) | Cell Signaling Technology<br>Cat# 9662,<br>RRID: AB_331439 | 1:1000 | WB |
| Caspase 3 (Cleaved) | Cell Signaling Technology<br>Cat# 9961,<br>RRID: AB_2341188 | 1:1000 | WB |
| CTGF | Santa Cruz Biotechnology<br>Cat# sc-365970,<br>RRID: AB_10917259 | 1:500 | WB |
| CYR61 | Santa Cruz Biotechnology<br>Cat# sc-374129,<br>RRID: | 1:500 | WB |
| DDR1 | Cell Signaling Technology<br>Cat# 5883,<br>RRID: AB_10947399 | 1:1000 | WB |
| Phospho-DDR1 (Tyr792) | Cell Signaling Technology<br>Cat# 11994,<br>RRID: AB_2797793 | 1:1000<br>1:200 | WB<br>IF |
| DDR2 | Cell Signaling Technology<br>Cat# 12133, | 1:1000 | WB |

| | | | |
|---|---|---|---|
| | RRID: AB_2797825 | | |
| Phospho-DDR1 (Tyr796)/DDR2 (Tyr740) | R&D Systems<br>Cat# MAB25382, | 1:1000<br>1:200 | WB<br>IF |
| E2F1 | Cell Signaling Technology<br>Cat# 3742,<br>RRID: AB_2096936 | 1:1000 | WB |
| ERK2 | Santa Cruz Biotechnology<br>Cat# sc-1647,<br>RRID: AB_627547 | 1:500 | WB |
| Phospho-ERK1/2 (Thr202/Tyr204) | Cell Signaling Technology<br>Cat# 9101,<br>RRID: AB_331646 | 1:1000 | WB |
| HSP60 | Santa Cruz Biotechnology<br>Cat# sc-59567,<br>RRID: AB_783870 | 1:500 | WB |
| HSP90 | Santa Cruz Biotechnology<br>Cat# sc-13119,<br>RRID: AB_675659 | 1:1000 | WB |
| Ki67 | Abcam Cat# ab-16667,<br>RRID: AB_302459 | 1:200 | IF |
| MITF | Sigma Cat# HPA003259,<br>RRID: AB_1079381 | 1:1000 | WB |
| MLC2 | Cell Signaling Technology<br>Cat# 8505,<br>RRID: AB_2728760 | 1:1000 | WB |
| Phospho-MLC2 (Ser19) | Cell Signaling Technology<br>Cat# 3671,<br>RRID: AB_330248 | 1:1000 | WB |
| Rb | Cell Signaling Technology<br>Cat# 9309,<br>RRID: AB_823629 | 1:1000 | WB |

| | | | |
|---|---|---|---|
| Phospho-Rb (Ser807/Ser811) | Cell Signaling Technology<br>Cat# 9308,<br>RRID: AB_331472 | 1:1000 | WB |
| Phospho-Tyrosine<br>(Tyr-100) | Cell Signaling Technology<br>Cat# 9411,<br>RRID: AB_331228 | 1:800 | IF |
| YAP | Cell Signaling Technology<br>Cat# 14074,<br>RRID: AB_2650491 | 1:1000<br>1:100 | WB<br>IF |
| ZEB1 | Santa Cruz Biotechnology<br>Cat# sc-25388,<br>RRID: AB_2217979 | 1:500 | WB |
| Anti-mouse IgG-HRP | Cell Signaling Technology<br>Cat# 7076,<br>RRID: AB_330924 | 1:2000 | WB |
| Anti-rabbit IgG-HRP | Cell Signaling Technology<br>Cat# 7074,<br>RRID: AB_2099233 | 1:2000 | WB |
| Goat anti-rabbit AF488 | ThermoFischer<br>Cat# A11034,<br>RRID: AB_2576217 | 1:200 | IF |
| Goat anti-mouse AF488 | ThermoFischer<br>Cat# A11029,<br>RRID: AB_2534088 | 1:200 | IF |
| Goat anti-rabbit AF594 | ThermoFischer<br>Cat# A11012,<br>RRID: AB_141359 | 1:200 | IF |
| Goat anti-mouse AF594 | ThermoFischer<br>Cat# A11005,<br>RRID: AB_141372 | 1:200 | IF |

WB : Western Blotting, IF : Immunofluorescence

**Supplementary Table 3**: List of qPCR primers

| Name | Forward sequence (5'->3') | Reverse sequence (5'->3') |
|---|---|---|
| AMOTL2 | GCGACTGTCAGAACAACTGC | GCACCTTTAACCTGCTTTCCA |
| AXL | GTGGGCAACCCAGGGAATATC | GTACTGTCCCGTGTCGGAAAG |
| CTGF | ACCGACTGGAAGACACGTTTG | CCAGGTCAGCTTCGCAAGG |
| CYR61 | TGAAGCGGCTCCCTGTTTT | CGGGTTTCTTTCACAAGGCG |
| CDH2 | CCACGCCGAGCCCCAGTAT | GGCCCCCAGTCGTTCAGGTAAT |
| CTNNB1 | CCGCAAATCATGCACCTTT | ATTTCTTCCATGCGGACCC |
| DDR1 | ATGGAGCAACCACAGCTTCTC | CTCAGCCGGTCAAACTCAAACT |
| DDR2 | AGTCAGTGGTCAGAGTCCACAGC | CAGGGCACCAGGCTCCATC |
| ERBB3 | GGTGATGGGGAACCTTGAGAT | CTGTCACTTCTCGAATCCACTG |
| HPRT | AGATGGTCAAGGTCGCAAGCT | AGTCAAGGGCATATCCTACAACAAAC |
| MITF | CATTGTTATGCTGGAAATGCTAGAA | GGCTTGCTGTATGTGGTACTTGG |
| MLANA | GCTCACTTCATCTATGGTTACCC | GACTCCCAGGATCACTGTCAG |
| NGFR | CCTACGGCTACTACCAGGATG | CACACGGTGTTCTGCTTGT |
| RPL32 | CCTTGTGAAGCCCAAGATCG | TGCCGGATGAACTTCTTGGT |
| Serpine1 | CCGCCTCTTCCACAAAATCA | CAGTTCCAGGATGTCGTAGTAATGG |
| SOX9 | CCCAACGCCATCTTCAAGG | GAGACCGTCCCTGCTGTGTT |
| SOX10 | CCTTCATGGTGTGGGCTC | CGCTTGTCACTTTCGTTCAG |
| Wnt5a | CAATGTCTTCCAAGTTCTTCCTAGTG | CATACCTAGCGACCACCAAGAAT |
| YAP1 | GCTACAGTGTCCCTCGAACC | CCGGTGCATGTGTCTCCTTA |
| ZEB1 | AGCGCTTCTCACACTCTGGGTCTT | CTGCCTCTTCCTGCTCTGTGC |
| ZEB2 | GCGCAGTGACACAGCCATTATTTA | CTTGTAGCCCCGGTCGCAGTA |

### Supplementary Figure 1

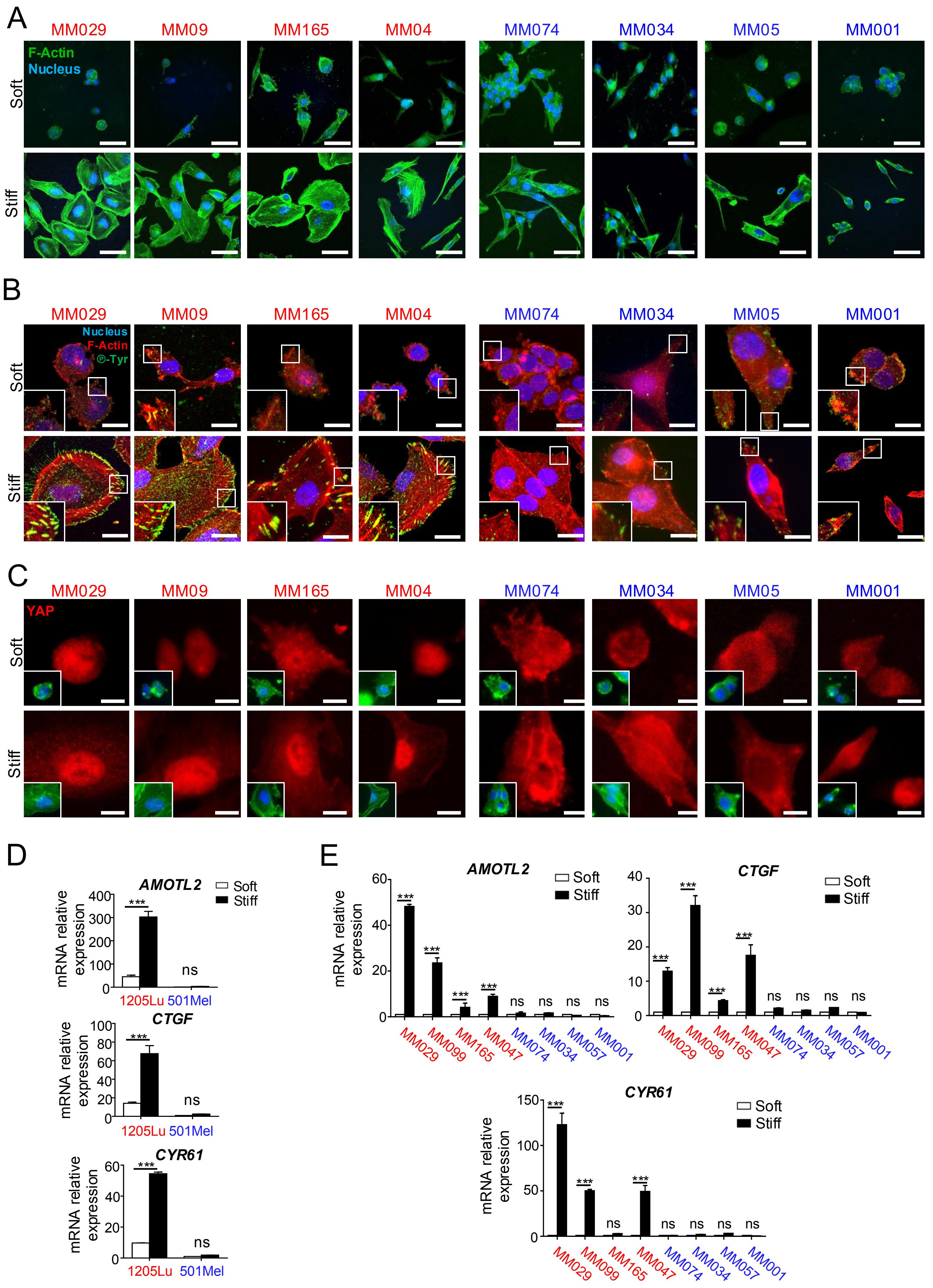

### Supplementary Figure 2

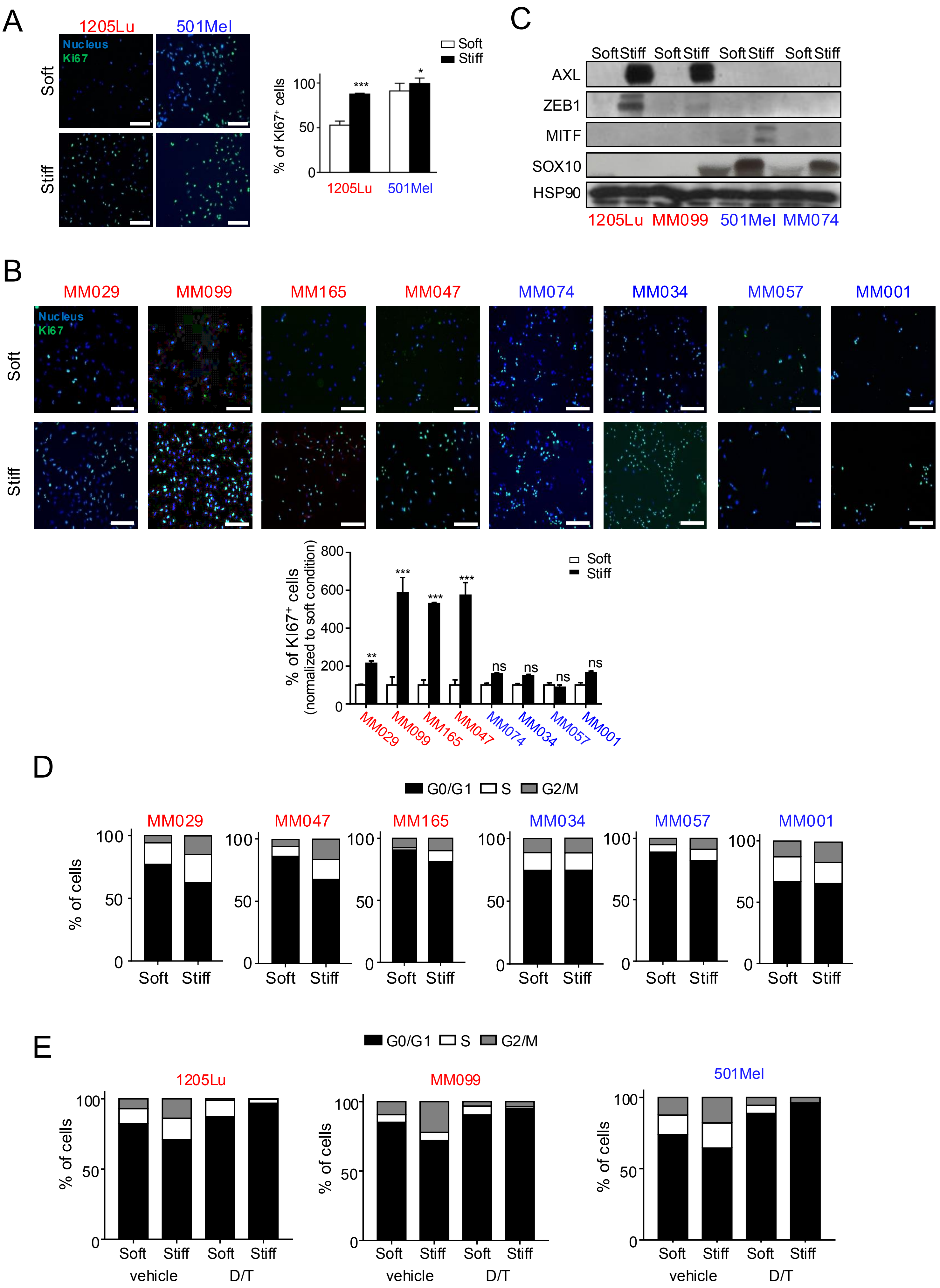

### Supplementary Figure 3

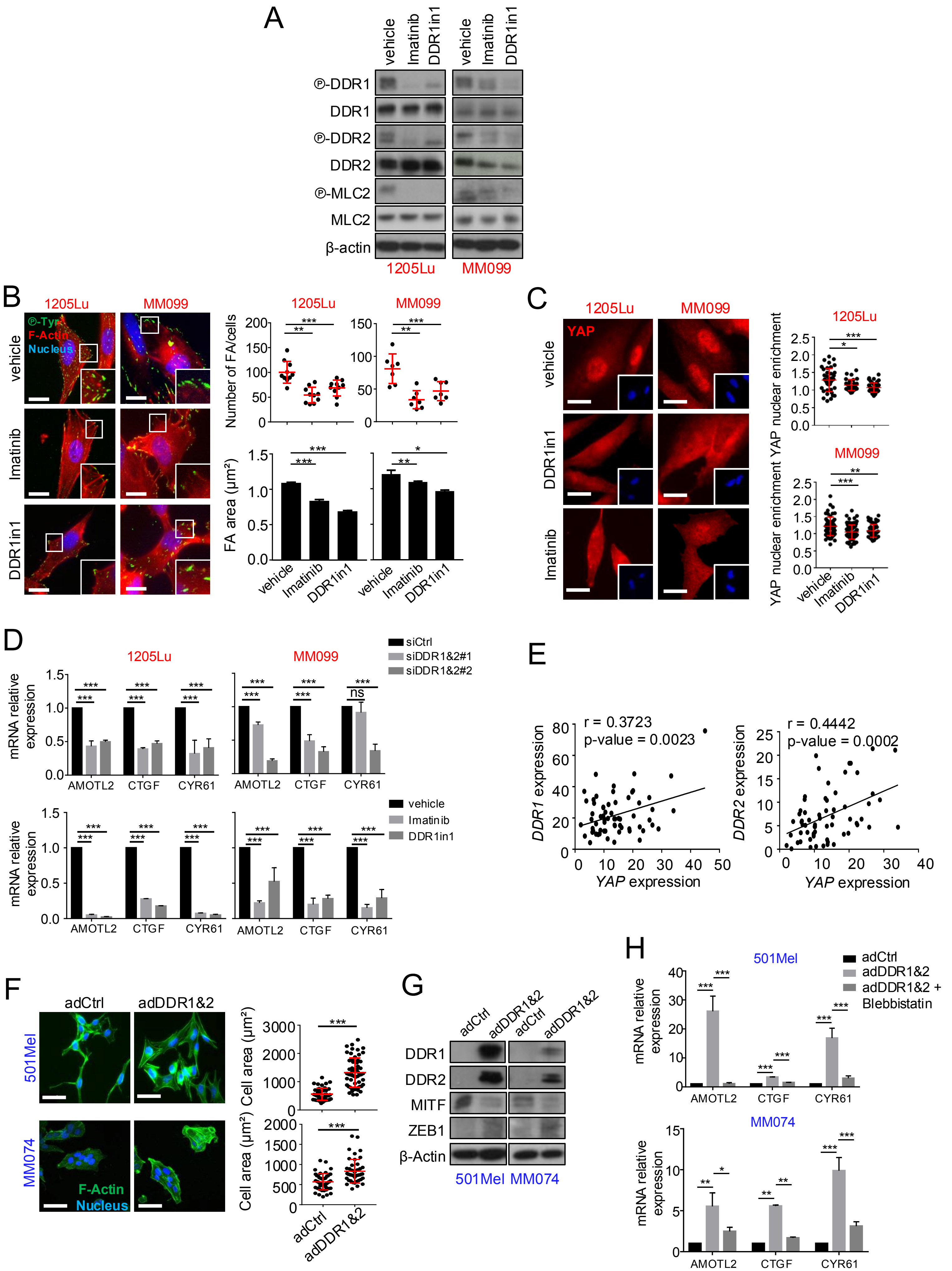

### Supplementary Figure 4

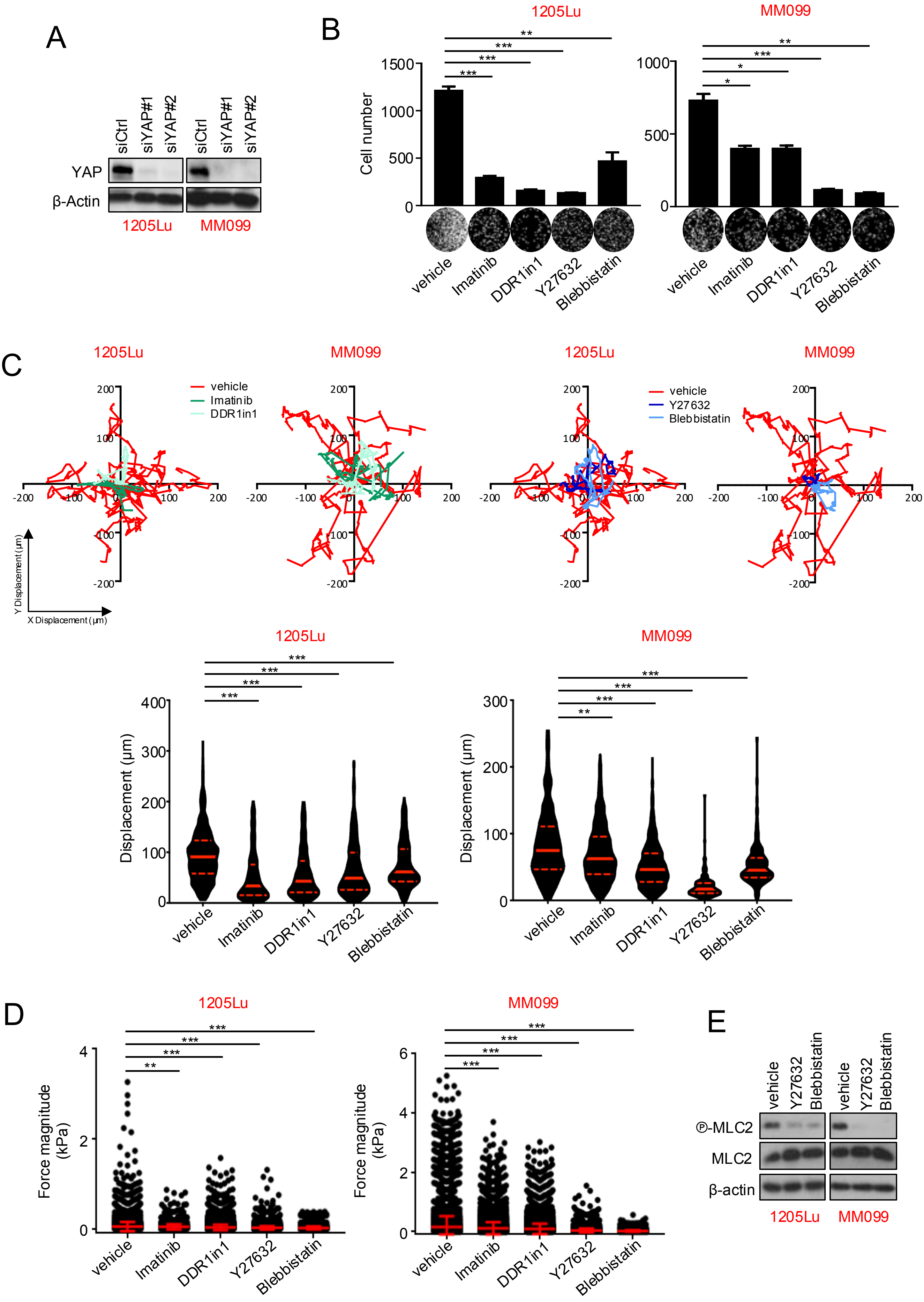

### Supplementary Figure 5

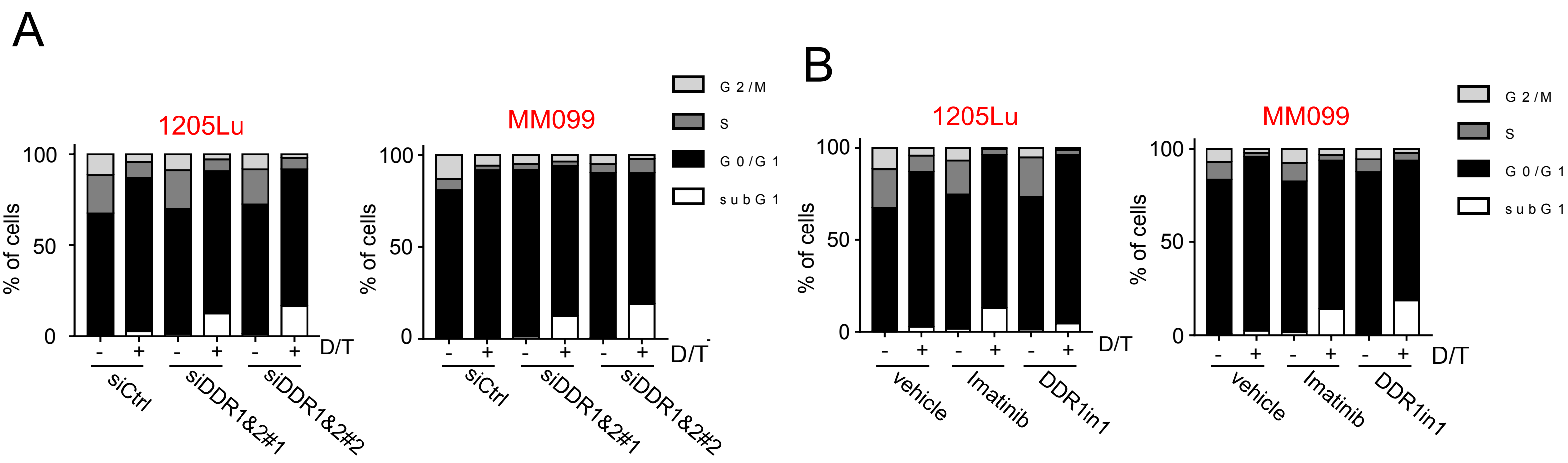
